# Head-to-head organized segmental paralogs *AtOFP2* and *AtOFP17* exhibit differential, spatio-temporal partitioning of function, and negative regulation of multiple developmental traits including seed-yield and root architecture

**DOI:** 10.64898/2026.06.30.735610

**Authors:** Nishu Chahar, Ekta Pokhriyal, Shobha Yadav, Bijie Ren, Meenakshi Dangwal, Sandip Das

## Abstract

Ovate Family Proteins (OFPs) are a class of plant-specific, negative nuclear transcriptional regulators characterized by conserved C-terminal OVATE domain. This study on comparative functional characterization of two head-to-head arranged OFPs – *AtOFP2* (Ovate-OFP with full ovate domain) and *AtOFP17* (Ovate-Like OFP with partial ovate domain) provides critical insight into how structural variations in ovate domain leads to functional divergence. Detailed phenotypic analysis of 28 physical and physiological traits of loss- and gain-of-function mutants revealed that both genes act as broad, pleotropic repressors of plant growth and development. Removal of repression in knock-down mutants of both genes exhibited reduced duration of seed dormancy, faster rate of germination and growth, bigger plants and significantly higher seed yield. In contrast, constitutive over-expression showed a generalized repressive nature of both genes, with nuanced differences for fine tuning of specific traits. For example, both genes showed antagonistic behaviours on root hair architecture. *AtOFP2* act as a strong repressor of root hair development whereas *AtOFP17* is a stronger repressor of hypocotyl and root cell architecture. *AtOFP17* owing to partial ovate domain exerts a mild level of repression throughout life span as indicated by smaller plants and lesser yield in knock-down *AtOFP17* mutants. On the contrary, *AtOFP2* exerted a much stronger repressor effect in which > 90% over-expression mutants died at the juvenile stage ; the survival of remaining 10% is probably owing to activation of dosage-dependent feedback loop mechanism as indicated by normal growth of mature plants, and is also evident by transcriptome data. Transcriptome analysis of roots of 7-day old seedling of knock-down and over-expression mutants of *AtOFP2* showed downregulation of *OFP2* in over-expressed mutants. However, severely stunted phenotype indicated presence of stable OFP2 protein to exert effects. Analysis of DEGs in *OFP2* mutants revealed that it acts as an important regulator working at intersection of hormonal signalling affecting critical genes required for auxin, cytokinin, GA, BR and ABA functioning. Perturbations across hormonal signalling pathways affects cell wall remodelling factors such as EXPANSINS, Xyloglucan hydrolases (XTHs) and cellulose synthases (CSLs) causing overall stunted growth; and epidermal patterning genes such as *WER, GL1, EGL3, TTG1* leading to severely reduced root length and root hairs. Significantly, functional analysis of this master regulator highlighted a significant economic potential. Knockdown of both these genes relieves their natural repression on reproductive traits, leading to longer siliques, bigger and heavier seeds, and substantially increased overall seed yield, positioning *AtOFP2* and *AtOFP17* as highly valuable targets for agricultural crop improvement.

## Introduction

Ovate Family Proteins (OFPs), characterized by conserved C-terminal OVATE domain are a novel class of plant-specific transcriptional regulators that play essential, yet undiscovered roles in plant development, growth and architecture (Wang et al, 2016). The ovate locus was identified more than a century ago, initially as “*pr”* locus and later termed as “*ovate”* or “*o”* locus responsible for pear shape fruit of tomato (Hedrick and Booth 1907; Prince and Drinkard 1908). Subsequently, *OVATE* locus was mapped on chromosome 2 of tomato as a major QTL (Ku et al., 1999), and the locus was isolated through map-based cloning (Liu et al., 2002). *OFPs* are now identified as a multi-gene family from *Marchantia*, *Physcomitrella*, *Selaginella* and *Sphagnum* (Dangwal and Das, 2018), lineages of embryophyta (Chahar et al., 2021, 2023), *A. thaliana* (Liu et al., 2014), *Oryza sativa* (Yu et al., 2015), to name a few.

The founder members were identified as determinants of fruit-shape (Tanksley, 2004), and characterized as nuclear localized transcriptional repressors (Hackbusch et al, 2005; Wang et al, 2007). Studies have now associated OVATE gene with regulating fruit shape not only in tomato (Rodriguez et al., 2011), but also in melon (Ma et al, 2022), *Citrus maxima (CitOFP19*; Wu et al., 2022); pepper (Tsaballa et al, 2011), radish (Wang et al, 2020), to name a few. Other investigations studies have established diverse roles of members of OFP family such as in vasculature development (Schmitz et al., 2015); ovule development (Pagnussat et al., 2007); secondary cell wall formation (Li et al., 2011); hormone mediated regulation of cell size (Wang et al., 2007) and fruit ripening (Liu et al., 2015), in different species, highlighting their importance as key growth regulators.

Mechanistically, *OFP1* and *OFP4* have been shown to form complexes with *BLH6* (BEL-Like Homeodomain 6) and *KNAT7* to modulate secondary cell wall formation (Li et al. 2011; Liu and Douglas, 2015). They also regulate cell elongation by repressing gibberellin synthesis through interaction with the *GA20ox1* promoter in *Arabidopsis*, pepper, and rice (Wang et al., 2007; Wang et al., 2010; Tsaballa et al., 2012; Schmitz et al., 2015). Over-expression of *OFPs* in *A. thaliana* results in stunted growth, altered organ shape and patterning due to negative regulation of cell division and cell elongation (Hackbusch et al, 2005; Zhu et al, 2024). For ex. over-expression of *AtOFP2* leads to kidney shaped cotyledons and curled leaves (Wang et al., 2011); *AtOFP2* and *AtOFP5* interact with TONNEAU2 (TON2), a major regulator of microtubule arrangement which direct their orientation in response to light and brassinosteroids (Zhang et al., 2020).

In tomato, *SlOFP20* interacts with the TRM family to produce the TTP complex (TON1, TRM, and PP2A), which is essential for microtubule array organisation, cell division pattern regulation, and, as a result, organ shape (Wang et al, 2020). In rice, *OsOFP8* has been shown to influence brassinosteroid signalling pathway through interacting with GSK3-like kinase, and regulate plant architecture via controlling inclination angle of lamina joint (Yang et al, 2016). A regulatory network of F-Box protein FBX206, *OsOFP8* and *OsOFP19* regulate brassinosteroid signalling to control grain size and yield in Rice (Sun et al, 2023). *OFPs* also contribute to stress responses like *OFP1* in *Prunus persica* interacts with *PpZFHD1* to confer salt tolerance (Tan et al., 2021), while *PpOFP1*in *Populus trichocarpa* and *TaOFP29a* in wheat are associated with drought tolerance (Wang et al., 2020; Wang et al., 2021). A survey of literature and comparative functional analysis suggests conserved nature of the sequence, and function of OVATE domains (Liu et al, 2014; Dangwal and Das, 2018; Chahar et al, 2023, Monforte et al. 2014; Wu et al., 2018; Colle et al. 2017).

Dissection of mode of action underscored the multifaceted role of *OFPs* in managing cross-talk between hormonal signals and their capacity to fine-tune plant developmental outcomes. Collectively, these studies demonstrate versatile and dynamic participation of *OFPs* in regulating plant growth and development through interactions within various regulatory pathways. Ovate Family Proteins (OFPs) are thus emerging as central components in complex regulatory networks integrating various hormonal and signalling pathways to influence plant development and architecture. Inspite of their near-universal presence across embryophyta as a multi-gene family, major knowledge gaps exist. For instance (i) only a handful of genes in selective species have been only partially functionally characterized and their mode and mechanism of actions are not fully understood; (ii) the full spectrum of plant developmental regulation via *OFPs* is yet to be unravelled, (iii) the functional outcome of structural variance of complete and the partial OVATE domain is not investigated and understood.

In *A. thaliana,* out of the 19 members of OFP-family (Liu et al., 2014), we reported that *AtOFP2-AtOFP17*, and *AtOFP4-AtOFP20* are present in head-to-head organization on chromosome 2 and 1, respectively (Chahar et al 2023). *AtOFP2* contains a complete OVATE domain (annotated as OVATE-OFP) with two motifs at C-terminal and a predicted DNA Binding Domain (DBD) at N- terminal, whereas *AtOFP17* contains partial OVATE-domain (OVATE-like OFP) with only a single motif, and without the DBD at N- terminal (Chahar et al, 2023). Our previous studies uncovered a close linkage between sequence and structural variant of the OVATE domain, and its evolutionary trajectory across the embryophyta, albeit without any insights into functional significance (Dangwal and Das 2018; Chahar et al., 2021, 2023). The presence of complete and partial domain structure in *AtOFP2* and *AtOFP17,* respectively, will permit testing a direct link between protein structure and function. The presence of both complete and partial OVATE domains within the OFP family is not merely an evolutionary novelty but likely underlies functional divergence.

The present study was therefore designed to perform a comparative functional characterization of *AtOFP2* and *AtOFP17* with the objectives of (i) to understand the effect of complete and partial-OVATE domain on plant development, (ii) perform functional analysis of *AtOFP17* which has not been reported yet, (iii) to comparative analyse the roles of *AtOFP2* and *AtOFP17* in plant growth and development using gain- and loss-of-function mutants, and (iv) to gain molecular insights into *OFP2* regulated root system architecture.

## Materials and methods

### Plant material, growth conditions

For all experiments, *Arabidopsis thaliana* ecotype Col-0 was used. Plants were grown in growth chamber maintained at 21 ± 2°C, 65% humidity, 16h-light and 8h-dark photoperiod cycle with 120-150 µmol/m^2^/sec light intensity (Weigel and Glazebrook, 2002).

### Generation of gain- and loss-of-function mutants

Transgenic lines were generated using Gateway™ technology for gain-of-function mutants. The CDS of *AtOFP2* and *AtOFP17* were PCR amplified (Supplementary table 1) and cloned into the pENTR/D-TOPO Entry vector (Thermo Fisher Scientific). For directional cloning, 5’-CACC-3’ was added at the 5’ end of forward primer. BP and LR recombination reactions were performed between pENTR/D-TOPO-*AtOFP2/AtOFP17* constructs and the destination vector pGWB441, which contains a 35S CaMV promoter and a C-terminal eYFP tag (Nakagawa et al., 2007a) to generate constructs for ectopic expression (Supplementary figure 1).

For loss-of-function mutants, full length CDS sequences of *AtOFP2* and *AtOFP17* were retrieved from TAIR and submitted to Web microRNA designer (http://wmd3.weigelworld.org/cgi-bin/webapp.cgi?page=Designer;project=stdwmd) server to design locus-specific artificial microRNAs. The target gene was compared against “Araport11 genes 201606 cDNA” database using the criterion of “minimum number of included targets = 1”, and “accepted off-targets = 0”. The most suitable artificial microRNA was selected based after manual examination of hybridization energy, mismatch positions, and target position (Supplementary figures 2, 3A-C). SOEing-PCR primers for artificial microRNA sequence in pRS300 backbone was designed, and the amplicon was engineered as described into pCHF3, and sequence verified (Schwab et al. 2006; Fankhauser et al., 1999; Supplementary figure 2, 3) prior to transforming into *Agrobacterium tumefaciens* strain GV3101.

*A. thaliana* Col-0 plants were transformed using floral dip method (Clough and Bent, 1998). Transformed plants were selected on kanamycin, and T1 plants were PCR genotyped. Plants were selfed to advance the generation and expression levels of both genes were confirmed in T2 plants via semi-quantitative PCR (Supplementary Figure 3D- E).

### Semi-quantitative RT-PCR

RNA was extracted from 15-day old seedlings at T2 generation using TRIzol™ (Thermo Fisher Scientific). cDNA synthesis was done using iScript™ kit (Bio-Rad; USA) and actin (*At3G18780*) was used as internal control for of transcript levels (Anand et al. 2023). Reaction conditions for first strand synthesis was 25⁰C for 5-mins; 46⁰C for 20 mins, and 96⁰C for 1 min. Semi-quantitative PCR was performed for 28-cycles using locus specific primers at 94⁰C, Tm-dependent annealing and 72⁰C for 20 secs each.

### Dormancy test

Dormancy in seeds of WT, *amiR*-*AtOFP2/AtOFP17*, and, *oex*-*AtOFP2/AtOFP17* was evaluated in the form of percent germination of non-stratified seeds along a defined time-period (Koornneef et al., 2002; Gubler et al., 2005). 100 seeds were uniformly sprinkled on ½ strength MS-Agar plates and directly kept in growth chamber at 22^0^C without stratification. On the first day, the seeds were exposed to 2-hours of light to initiate germination followed by 8h-dark period. For all subsequent days, a 16h-8h light-dark cycle was followed. Number of seeds germinated were observed and photographed using Stemi-305 Stereozoom microscope (Carl-Zeiss, Switzerland). Seeds where radicle emerged after penetrating the seed coat were considered as germinated.

### Seed morphology and physiology

To investigate phenotypic differences, general characters such as seed size, colour, weight, and, physiological properties like mucilage production, permeability of seed coat and generation of proanthocynidines were assessed through electron microscopy, and through histochemical staining with Ruthenium red, DMACA, and Tetrazolium salt.

### Scanning electron microscopy

Seeds were oven dried at 37^0^C before analysis for seed coat pattern. Seeds were subjected to critical point drying for 30 mins in the presence of liquid CO2 and then mounted on stub and coated with gold in sputter coater. The samples were subsequently viewed under 10kV and 20kV and photographed using Scanning Electron Microscope (JEOL, 730 JSM-6610LV).

### Ruthenium red staining

Seeds were placed in a 1.5 ml microcentrifuge tube (MCT) and 500 µl water was added to completely submerge the seeds. The contents were vigorously shaken in an open rotary shaker at 400 rpm for 1 hr for proper hydration and release of mucilage. After 1-hour incubation, water was replaced with 0.01% ruthenium red dye for 2 hours (McFarlane et al., 2014). Seeds were then washed with water, observed and photographed using Stemi-305 Stereozoom microscope (Carl-Zeiss, Switzerland).

### DMACA (Dimethylamino-cinnamaldehyde) staining

Seeds were directly placed in DMACA solution (0.2% DMACA stain in 3M-HCL/50% Methanol). After 5-days, seeds were washed in 70% ethanol, observed and photographed using Stemi-305 Stereozoom microscope (Li et al., 1996; Carl-Zeiss, Switzerland).

### Tetrazolium salt staining

Seeds were directly placed in tetrazolium staining solution (0.1 gm Triphenyl tetrazolium chloride (2,3,5-triphenyl-2H-tetrazolium chloride) in 10 ml sterile water) for 24 to 36 hours. After staining, seeds were washed with water and photographed using Stemi-305 Stereozoom microscope (Carl-Zeiss, Switzerland).

### Statistical analysis

ANOVA, Tukey’s test, and regression analysis were done in GraphPad Prism 9.3.1 (https://www.graphpad.com/scientific-software/prism/; GraphPad Software, Inc., San Diego, California, USA; Mavrevski et al., 2018).

### RNA-sequencing and data analysis of *AtOFP2* seedling roots

Roots of 7-day old seeding were used for RNA extraction; RNA sequencing libraries were constructed for WT, *oex-ofp2* and *amiR-ofp2* in triplicate for each sample, and sequenced on Illumina NovaSeq 6000 platform. For each sample 35-40 million reads were generated. Raw RNA-seq reads were quality-checked and trimmed using Trimmomatic (v 0.39) to remove low-quality bases and adapter sequences (R-code in supplementary data file). High-quality trimmed reads were then aligned to the reference *A. thaliana* genome TAIR 10 in STAR 2.7.10 (Spliced Transcripts Alignment to a Reference; Dobin et al. 2013). The sorted BAM files generated from STAR were used for gene-level quantification with HTSeq-count (v 0.12.4). The GTF file corresponding to the reference genome was provided to assign reads to annotated genes. Raw gene counts, generated from HTSeq-count, were imported into R along with the corresponding sample metadata. To improve the reliability of statistical tests, genes with very low counts across all samples were filtered out. The differential expression analysis was performed using DESeq2 core method, which models the counts using a negative binomial distribution and estimates dispersion for each gene. DESeq2 then applies a generalized linear model (GLM) to test for differences in gene expression between conditions. Genes with a log2 fold change greater than 1 (log2FC > 1) were selected and analysed for GO enrichment using agriGO 2.0 platform (Tian et al. 2017; https://systemsbiology.cau.edu.cn/agriGOv2/). Venn diagrams were generated using Venny 2.1 (https://bioinfogp.cnb.csic.es/tools/venny/index2.0.2.html; Oliveros 2007-2015).

### Real-time qRT-PCR

Total RNA was isolated from the roots of 7-days old seedlings of *A. thaliana* Col-0 (WT), *amiR-ofp2* and *oex-ofp2* lines using Trizol Reagent (Thermo Fisher Scientific); First strand cDNA synthesis was performed using Verso cDNA synthesis kit (Thermo Fisher Scientific). Real time qRT-PCR was used to perform the relative quantification of the selected candidate genes using KAPA SYBR FAST (KAPA biosystems; qTower by Analytiykjena). The CDS sequence of the selected candidate genes was used to manually design primers towards the 3’end. Each 10 μL reaction mixture contained 5 μL of 2×SYBR Premix, 0.5 μL of first strand synthesis reaction, 0.5 μL of each primer (2 μM), and 3.5 μL of nuclease-free double distilled water.

The qPCR cycling conditions were as follows: 95°C for 30 s; followed by 40 cycles of 95°C for 20 s, 55°C annealing temperature for 20 s and extension at 72°C for 30 s in PCR plates. Actin was used as an internal control to normalise the variance between the samples. The data was visualised using qPCR Soft 4.1 and the data was exported to the excel sheet. The level of transcript and the fold change was analysed using 2-ΔΔCt method (Livak and Schmittgen, 2001).

## Results

### Phenotypic and physiological analysis reveal repressive and pleotropic effects of *AtOFP2* and *AtOFP17*

To investigate the roles of *AtOFP2* and *AtOFP17* in *A. thaliana* development, gain-of-function and loss-of-function mutant lines were generated using constitutive over-expression (*Oex*) and amiRNA-based knock-down (*amiR*) constructs. Transcript levels of target loci were confirmed at T2 generation. Significant increase in transcript level of *AtOFP2* and *AtOFP17* in respective over-expression, and decrease in knock-down lines were observed compared to WT (Supplementary figure 3D, E). Evaluation of phenotypic trait was done at T3 generation with 10-15 progenies of atleast three independent transgenic lines. As nothing was previously reported about the phenotypes affected by mis-expression of *AtOFP2* and *AtOFP17*, detailed phenotypic assessments across the entire life cycle, from seed germination to final plant yield was undertaken. Initial screens revealed similar phenotypes for both genes but detailed analysis throughout life cycle indicated towards difference in the penetration of effects of mutation for the two genes.

## Seedling traits

### Cotyledonary leaf area

The area of cotyledonary leaves in 7-day-old seedlings showed significant variation across genotypes. 15 individual T3 plants per genotype were assessed (Supplementary Table S2a, S2d). In *AtOFP2*, maximum cotyledonary leaf area was observed in *amiR-ofp2* (av. 5.132 ± 0.227 mm²), followed by WT (av. 3.016 ± 0.219 mm²), and the smallest in *oex-ofp2* (av. 1.386 ± 0.126 mm²; Figure 1A, 1F). One-way ANOVA revealed significant difference within groups in size of cotyledon area with a F-value of 1368 and P-value <0.001 (Fcrit- 3.220; Supplementary Table S2b). To find out the significance of variation between groups in a pair wise manner Tukey’s test was performed. Q-value for *amiR-ofp2* vs. WT, *amiR-ofp2* vs. *oex-ofp2* and WT vs. *oex-ofp2* were 41.66, 73.76 and 32.1 respectively (Supplementary Table S2c). All these values were significantly higher than q.05 crit- 3.44 and q.01 crit- 4.35 making all the groups significantly different from each other (Supplementary Table S2c). The goodness-of-fit was found to be 0.9849 (R2 value; Supplementary Table S2b), and explains that the variation in size of cotyledons is genotype dependent.

**Figure 1:**
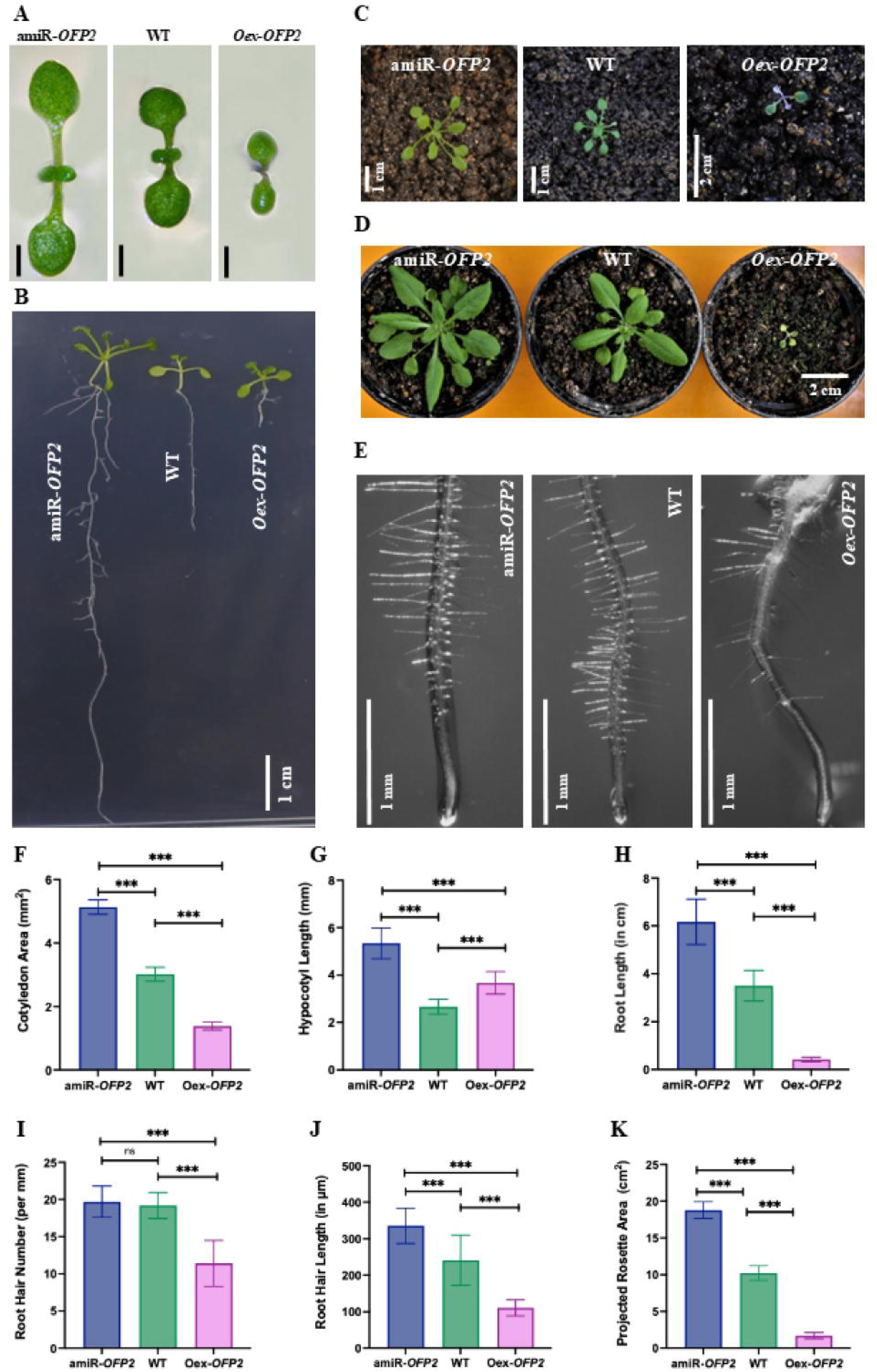
Comparison of seedling traits and rosette area between WT, knock-down (*amiR-OFP2*) and over-expression (*oex-OFP2*) plants. **A)** Cotyledonary leaves of 7-days old amiR-*OFP2*, WT and oex-*OFP2* seedlings. **B)** 15-days old seedlings exhibiting altered root morphology. **C, D)** 3-weeks old plants and 5-weeks old plants. **E)** Representative pictures of the phenotype of root hair length and root hair density. **F, G, H, I, J** and **K)** Graphical representation of statistical comparisons of cotyledon area, hypocotyl length, root length, root hair density, root hair length and projected rosette area, respectively. Bars in all graphs represent mean ± SD, ns = no significant difference, *p=0.033, **p=0.02, ***p<0.001.

Similarly, in *AtOFP17* lines, cotyledons with largest area were in *amiR-ofp17* (av. 4.326 ±0.375 mm²), followed by WT (av. 3.236 ±0.431 mm²), and the smallest cotyledons were observed in *oex-ofp17* seedlings (av. 2.202±0.31 mm² Figure 2A, 2F; Supplementary Table S2d). One way ANOVA revealed significant difference within groups in size of cotyledon area with a F-value of 119.2 at P-value <0.001 which is higher than Fcrit- 3.220 (Supplementary Table S2e). Q-value for *amiR-ofp17* vs. WT, *amiR-ofp17* vs. *oex-ofp17* and WT vs. *oex-ofp17* were 11.23, 21.89 and 10.66 respectively. All these values were significantly higher than q.05 crit- 3.44 and q.01 crit- 4.35 making all the groups significantly different from each other (Supplementary Table S2f). The goodness-of-fit was found to be 0.8509 revealing that along with genotype, other factors are also responsible for phenotype (Supplementary Table S2e).

**Figure 2:**
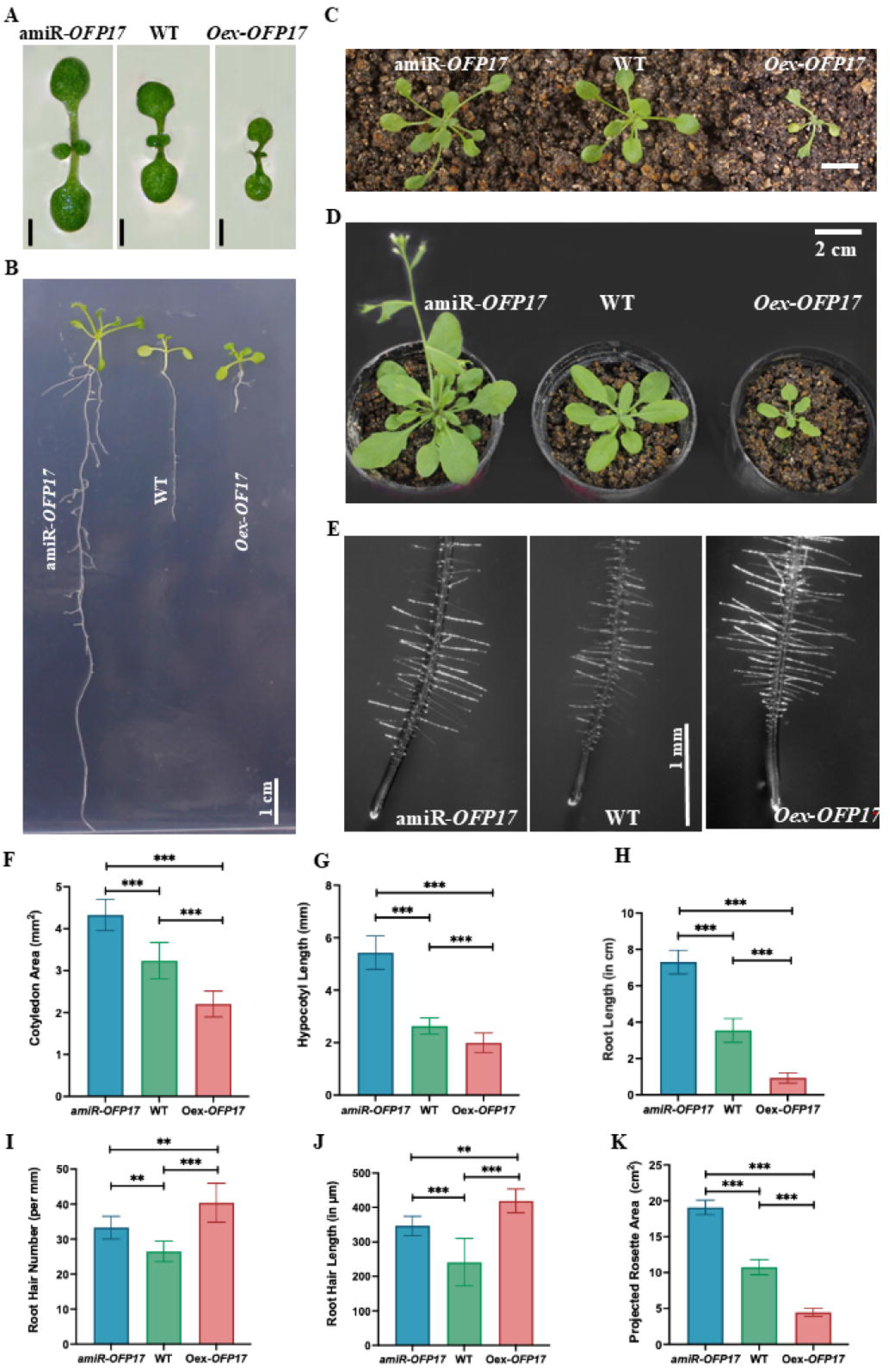
Comparison of seedling traits and rosette area between WT, knock-down (*amiR-OFP17*) and over-expression (*oex-OFP17*) plants. **A)** Cotyledonary leaves of 7-days old *amiR-OFP17*, WT and *oex-OFP17* seedlings. **B)** 15-days old seedlings exhibiting altered root morphology. **C, D)** 3-weeks old plants and 5-weeks old plants. **E)** Representative pictures of the phenotype of root hair length and root hair density. **F, G, H, I, J** and **K)** Graphical representation of statistical comparisons of cotyledon area, hypocotyl length, root length, root hair density, root hair length and projected rosette area, respectively. Bars in all graphs represent mean ± SD, ns = no significant difference, *p=0.033, **p=0.02, ***p<0.001.

### Hypocotyl length

The trait was measured in 15 individuals of T3 seedlings per genotype at 10-days after germination. In both *AtOFP2* and *AtOFP17*, knockdown lines exhibited the longest hypocotyls, followed by wild type (WT) with overexpression lines having the shortest (Figures 1B, 1G; 2B, 2G). For *AtOFP2, amiR-ofp2* seedlings showed an average hypocotyl length of 5.34 ± 0.65 mm compared to WT (av. 2.66 ± 0.31 mm) and *oex-ofp2* (av. 3.67 ± 0.47 mm; Supplementary Table S3a). For *AtOFP17*, average hypocotyl length in *amiR-ofp17* was 5.43 ± 0.64 mm, compared to 2.64 ± 0.31 mm in WT and 1.99 ± 0.37 mm in *oex-ofp17* (Supplementary Table S3d). One-way ANOVA indicated significant differences among genotypes for both genes (*AtOFP2*: F = 109.8, Fcrit = 3.22; *AtOFP17*: F = 235.3, Fcrit = 3.22; P < 0.001; Supplementary Table S3b, e). Tukey’s test confirmed that all pairwise comparisons were significant (q-values > 7.8 for *AtOFP2* and > 5.3 for *AtOFP17*, exceeding q₀.₀₅crit = 3.44 and q₀.₀₁crit = 4.35; Supplementary Table S3c, f). Goodness-of-fit was 0.84 for *AtOFP2* and 0.92 for *AtOFP17*, indicating a stronger genotype–phenotype relationship for *AtOFP17*, whereas additional factors may also influence the trait for *AtOFP2*.

### Primary root length

Measurements taken from 15-day-old seedlings showed a clear trend across both *AtOFP2* and *AtOFP17* genotypes: amiRNA lines had the longest roots, WT displayed intermediate lengths, and overexpression lines had markedly shorter roots (Figures 1B, 1H; 2B, 2H). In *AtOFP2*, *amiR-ofp2* root length averaged 6.17 ± 0.95 cm, more than double the WT length (3.50 ± 0.64 cm) and substantially longer than *oex-ofp2* (0.41 ± 0.09 cm; Supplementary Table S4a). A similar pattern emerged in *AtOFP17*, where *amiR-ofp17* root length measured 7.30 ± 0.66 cm compared with 3.54 ± 0.66 cm for WT and only 0.93 ± 0.28 cm for *oex-ofp17* (Supplementary Table S4d). Statistical analysis confirmed these differences were highly significant. One-way ANOVA produced F-values of 470.3 for *AtOFP2* and 816.2 for *AtOFP17*, well above the critical threshold (Fcrit = 3.12; P < 0.001; Supplementary Tables S4b, S4e). Tukey’s post hoc test separated all genotypes with large q-value differences (all > 20.1 for *AtOFP2* and > 23.2 for *AtOFP17*), surpassing both q₀.₀₅crit = 3.38 and q₀.₀₁crit = 4.25 (Supplementary Tables S4c, S4f). The high R² values (0.93 for both genes) indicate that root length variation is strongly associated with genotype.

### Root hair density

Root hair density was measured in the maturation zone of 15-day-old seedlings. In *AtOFP2* lines, *amiR-ofp2* and WT had similar hair density (19.7 ± 2.11 hairs/mm² and 19.2 ± 1.75 hairs/mm², respectively), and *oex-ofp2* had the lowest (11.4 ± 3.09 hairs/mm²; Figure 1E, 1I; Supplementary Table S5a). In contrast, *AtOFP17* showed an opposite trend, with *oex-ofp17* displaying the highest density (40.4 ± 5.56 hairs/mm²), followed by WT (26.5 ± 2.95 hairs/mm²) and *amiR-ofp17* (33.3 ± 3.23 hairs/mm²) (Figures 2E, 2I; Supplementary Table S5d).

ANOVA detected significant genotype effects for both genes (*AtOFP2*: F = 37.96; *AtOFP17*: F = 28.92; Fcrit = 3.35; P < 0.001; Table S5b, S5e). For *AtOFP2*, Tukey’s test showed no significant difference between *amiR-ofp2* and WT (q = 0.66), but both differed significantly from *oex-ofp2* (q = 10.99 and 10.32; q₀.₀₅crit = 3.51, q₀.₀₁crit = 4.49; Supplementary Table S5c). In *AtOFP17*, all genotypes were significantly different, with the largest separation between WT and *oex-ofp17* (q = 10.75; Supplementary Table S5f).

Interestingly, *amiR-ofp2* enhanced whereas *oex-ofp2* suppressed root hair density. In contrast in *AtOFP17 amiR-ofp17* suppressed whereas *oex-ofp2* enhanced root hair density exhibiting a reverse developmental pattern, suggesting these two *OFP* genes may have opposing regulatory roles in root hair development.

### Root hair length

Length of root hair was assessed in 15-day-old seedlings using 10 T3 plants per genotype and 30 hairs per plant. In *AtOFP2* lines, *amiR-ofp2* produced the longest hairs (335.5 ± 48.22 µm), WT was intermediate (241.2 ± 68.77 µm), and *oex-ofp2* the shortest (110.5 ± 21.99 µm; Figure 1E, 1J; Supplementary Table S6a). In *AtOFP17*, the trend reversed: *oex-ofp17* exhibited the longest hairs (419.1 ± 34.35 µm), followed by *amiR-ofp17* (346.5 ± 28.09 µm), with WT again at the low end (241.2 ± 68.77 µm) (Figures 2E, 2J; Supplementary Table S6d). ANOVA confirmed strong genotype effects (*AtOFP2*: F = 50.82; *AtOFP17*: F = 35.83; Fcrit = 3.35; P < 0.001; Supplementary Tables S6b, S6e). In *AtOFP2*, all pairwise comparisons were highly significant (q = 5.95–14.2; q₀.₀₅crit = 3.38; q₀.₀₁crit = 4.25; Supplementary Table S6c). In *AtOFP17*, *amiR-ofp17* differed strongly from both WT and *oex-ofp17* (q = 7.05 and 11.9), while WT vs. *oex-ofp17* was only marginally above the q₀.₀₁crit threshold (q = 4.85; Supplementary Table S6f). R² values were moderate (*AtOFP2*: 0.79; *AtOFP17*: ∼0.8), suggesting that additional factors beyond genotype contribute to root hair length variation.

Interestingly, *amiR-ofp2* enhanced whereas *oex-ofp2* suppressed length of root hair, whereas a different response was observed in reverse genetic mutants of *AtOFP17*. *oex-ofp17* produced longest root hairs followed by *amiR-ofp17*. This mirrors the pattern seen in root hair density and reinforces the possibility of opposing regulatory roles for *AtOFP2* and *AtOFP17* genes in root hair development.

### Root cell dimensions

Cell dimensions in the maturation zone of 15-day-old roots were quantified using images from 10 seedlings of T3 generation per genotype. In *AtOFP2* lines, *amiR-ofp2* cells were longer (174.17 ± 10.52 µm) than *oex-ofp2* (68.41 ± 4.65 µm) whereas WT has cells of intermediate length (137 ± 7.8 µm). Cell width followed a similar pattern, with *amiR-ofp2* cells wider (29.54 ± 4.86 µm) than *oex-ofp2* (14.48 ± 1.31 µm) and WT (18.10 ± 3.09 µm; Supplementary Figure 4A; Supplementary Table S7a). In *AtOFP17*, cells were longer (211.48 ± 8.74 µm) in *amiR-ofp17* than *oex*-*ofp17* (42.92 ± 3.84 µm) and WT (137 ± 7.8 µm). Cells were also wider in *amiR-ofp17* (35.95 ± 6.87 µm), followed by WT (18.10 ± 3.09 µm), while *oex-ofp17* cells were narrow (6.79 ± 1.07 µm; Supplementary Figure 5A; Supplementary Table S7f). ANOVA confirmed significant genotype effects for both traits (*AtOFP2*: F = 446.9 for length, 53.04 for width; *AtOFP17*: F = 1408 for length, 111.8 for width; Fcrit = 3.35; P < 0.001; Supplementary Tables S7b, d, g, i). Tukey’s test for *AtOFP2* showed strong separation of cell length among all genotypes (q = 14.62–41.67; Table S7c) and significant width differences except between WT and *oex-ofp2* (q = 3.35; Table S7e). For *AtOFP17*, all pairwise comparisons for both length and width were highly significant (length q = 33.05–74.88; width q = 8.13–20.97; Supplementary Tables S7h, j).

R² values indicated that variation in cell length is largely genotype-driven (*AtOFP2*: 0.97; *AtOFP17*: ∼0.97), whereas cell width showed lower fit (*AtOFP2*: 0.80), suggesting additional factors influence this trait.

### Potted plant traits

#### Projected rosette area

At the 5-week stage, amiRNA lines of both *OPF2* and *OFP17* exhibited the largest rosette areas. In *amiR-ofp2*, rosette area averaged 18.78 ± 1.22 cm², compared to 10.22 ± 0.98 cm² for WT and 1.70 ± 0.42 cm² for *oex-ofp2* (Figure 1D, 1K; Supplementary Table S8a). Rosette area in *amiR-ofp17* was 19.07 ± 1.02 cm², 10.74 ± 1.05 cm² in WT, and 4.88 ± 0.55 cm² in *oex-ofp17* (Figure 2D, 2K; Supplementary Table S8d). ANOVA detected highly significant genotype effects (*AtOFP2*: F = 912.7; *AtOFP17*: F = 650.7; Fcrit = 3.35; P < 0.001; Supplementary Tables S8b, S8e). Tukey’s test separated all groups with large q-values (*AtOFP2*: q = 30.3–60.4; *AtOFP17*: q = 21.9–50.9), well above q₀.₀₅crit = 3.44 and q₀.₀₁crit = 4.35 (Supplementary Table S8c, f). R² values were very high (*AtOFP2*: 0.99; *AtOFP17*: ∼0.98), indicating a strong genotype–phenotype relationship for rosette area in both genes.

### Number of rosette leaves

The number of rosette leaves at bolting stage in *amiR-ofp2* and WT plants was comparable (12.6 ±1.45 and 12.2 ±2.21, respectively), while *oex-ofp2* showed a significant increase (23.2 ± 2.12; Figure 3A – D; Supplementary Table S9a). Similarly, *amiR-ofp17* and WT showed similar leaf counts (12.8 ±1.61 and 13.26 ± 1.28, respectively), whereas *oex-ofp17* had a lower number (10 ± 1.13; Figure 4A – D; Supplementary Table S9d). ANOVA detected significant differences among genotypes (*AtOFP2*: F = 154.2; *AtOFP17*: F = 25.43; Fcrit = 3.22; P < 0.001; Supplementary Tables S9b, S9e). Tukey’s test showed no difference between *amiR-ofp2* and WT (q = 0.79) or between *amiR-ofp17* and WT (q = 1.33), while over-expression lines differed strongly from both knock-down and WT in each gene (*AtOFP2*: q = 21.1–21.9; *AtOFP17*: q = 7.99–9.32; q₀.₀₅crit = 3.51; q₀.₀₁crit = 4.49; Supplementary Table S9c, f). R² values were moderate (*AtOFP2*: 0.88; *AtOFP17*: 0.54), indicating that while genotype contributes significantly, other factors may also influence leaf number.

**Figure 3:**
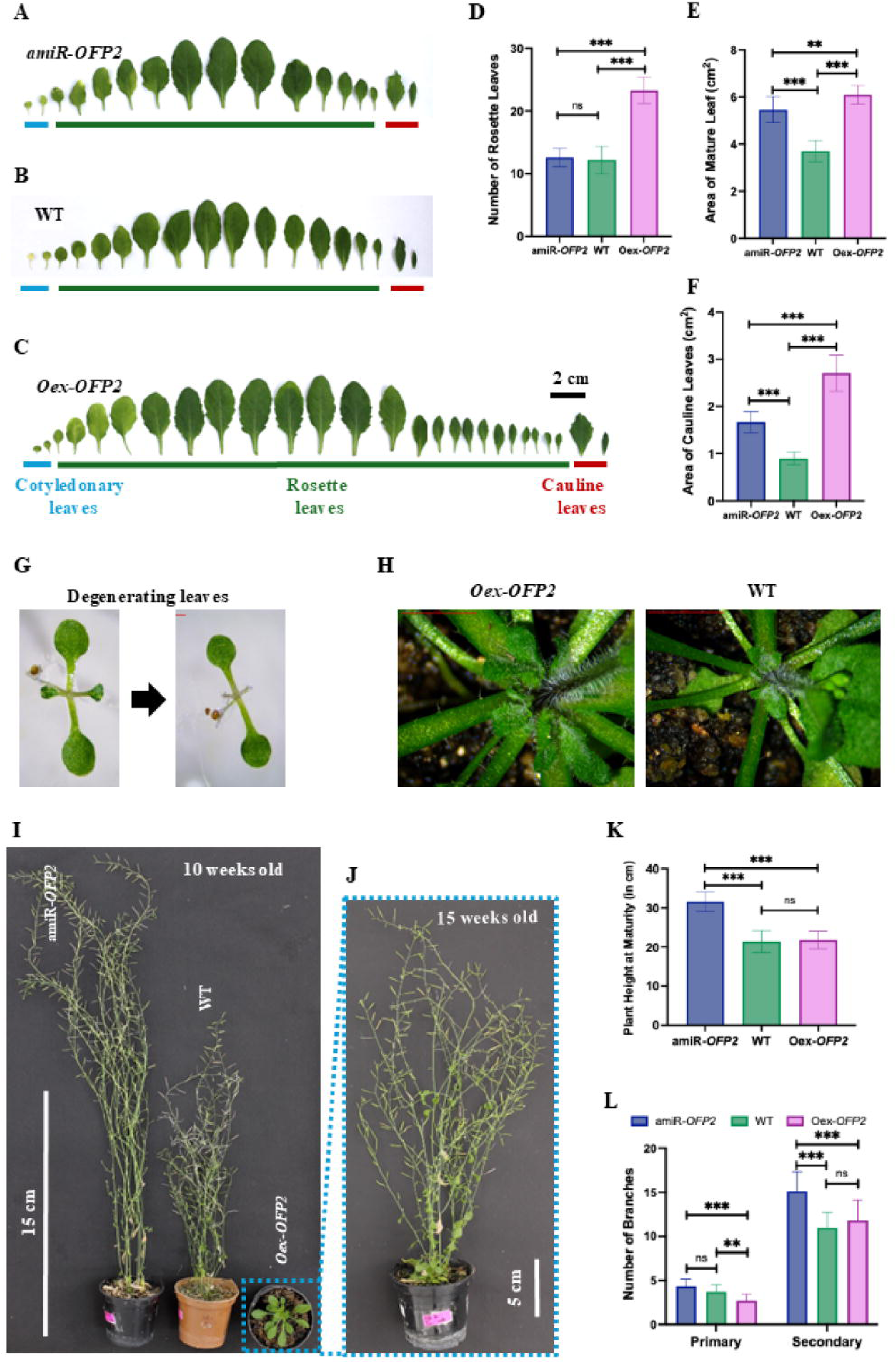
Comparison of mature plant phenotypes between WT, knock-down (*amiR-OFP2*) and over-expression (*oex-OFP2*) plants. **A, B, C)** Cotyledonary, rosette and cauline leaves of *amiR-OFP2*, WT and *oex-OFP2* plants, respectively. **D, E, F)** Graphical representation of statistical comparisons of number of rosette leaves, area of rosette leaves and area of cauline leaves, respectively. **G)** Degeneration of first true leaves in oex-*OFP2* plants. H) Whorl of leaves emerging from base of stem after bolting in *oex-ofp2* and pair of leaves from WT plants. **I, J)** 10-weeks old plants of *amiR-OFP2,* WT and *oex-OFP2;* and 15-weeks old plant of *AtOFP2*-oex respectively. **K, L)** Graphical representation of statistical comparisons of plant height at maturity; and primary and secondary branches, respectively. Bars in all graphs represent mean ± SD, ns = no significant difference, *p=0.033, **p=0.02, ***p<0.001.

**Figure 4:**
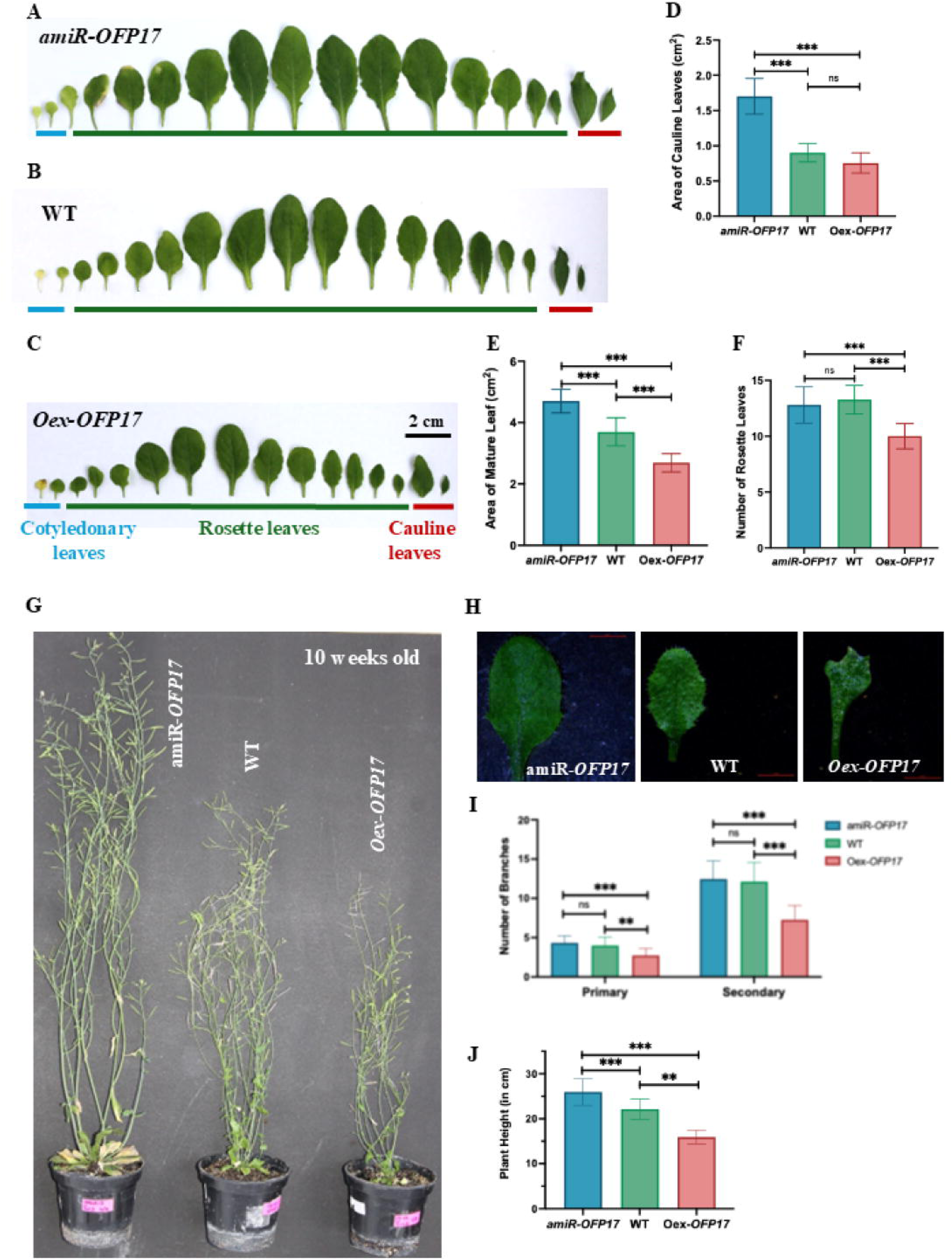
Comparison of mature plant phenotypes between WT, knock-down (*amiR-OFP17*) and over-expression (*oex-OFP17*) plants. **A, B, C)** Cotyledonary, rosette and cauline leaves of *amiR-OFP2*, WT and *oex-OFP17* plants, respectively. **D, E, F)** Graphical representation of statistical comparisons of area of cauline leaves, area of rosette leaves and number of rosette leaves, respectively. **G)** 10-weeks old plants of *amiR-OFP17*, WT and *oex-OFP17.* **H)** Morphology of leaf from potted seedling of *amiR-ofp17*, WT and *oex*- *OFP17* plants. **I, J)** Graphical representation of statistical comparisons of primary branches and secondary branches, and plant height, respectively. Bars in all graphs represent mean ± SD, ns = no significant difference, *p=0.033, **p=0.02, ***p<0.001.

### Area of rosette leaves and cauline leaves

At the rosette stage, *AtOFP2* lines showed the largest leaf area in *oex-ofp2* (6.09 ± 0.40 cm²), followed by *amiR-ofp2* (5.47 ± 0.55 cm²) and WT (3.70 ± 0.45 cm²; Figure 3A – C, E; Supplementary Table S10a). Contrary to this, leaves of *amiR-ofp17* were the largest (4.70 ± 0.38 cm²), followed by WT (3.70 ± 0.46 cm²), and smallest in *oex-ofp17* (2.69 ± 0.29 cm²) (Figure 4A – C, E; Supplementary Table S10d). ANOVA confirmed significant genotype effects (*AtOFP2*: F = 100.9; *AtOFP17*: F = 101.9; Fcrit = 3.22; P < 0.001; Supplementary Table S10b, e). Tukey’s test separated all *AtOFP2* genotypes (q = 5.07–19.37) and all *AtOFP17* genotypes (q = 10.02–20.19), well above q₀.₀₅crit = 3.44 and q₀.₀₁crit = 4.35 (Supplementary Tables S10c, S9f). R² values were moderate (*AtOFP2*: 0.83; *AtOFP17*: 0.83), suggesting additional factors contribute to rosette leaf area variation.

Cauline leaf area averaged 1.67 ± 0.22 cm² in *amiR-ofp2*, 0.90 ± 0.13 cm² in WT, and 2.70 ± 0.38 cm² in *oex-ofp2* (Figure 3A – C, F; Supplementary Table S11a). The corresponding values were 1.70 ± 0.25 cm² in *amiR-ofp17* and 0.75 ± 0.14 cm² in *oex-ofp17* (Figure 4A – C, F; Supplementary Table S11d). ANOVA again showed strong genotype effects (*AtOFP2*: F = 173.8; *AtOFP17*: F = 113.8; P < 0.001; Supplementary Table S11b, e). Tukey’s test separated all *AtOFP2* genotypes (q = 5.07–26.27) and identified significant differences for *AtOFP17* between amiRNA lines and both WT and oex (q = 16.76–19.81). WT vs. *oex-ofp17*, however, was not significant (q = 3.05; Figure 4D; Supplementary Table S11f). R² values were high for *AtOFP2* cauline leaves (0.89), indicating genotype as a major factor, while *AtOFP17* showed a slightly weaker fit (0.84), suggesting additional influences.

### Stem trichome density

Trichomes were counted over a 2 mm segment of the primary stem in ∼7-week-old plants. Both *amiR-ofp2* (13.6 ± 1.34) and *oex-ofp2* (13.3 ± 1.70) showed markedly higher trichome density than WT (5.1 ± 1.19). A similar pattern was observed in *AtOFP17*, with *amiR-ofp17* (12.8 ± 1.22) and *oex-ofp17* (12.8 ± 1.62) both exceeding WT (5.1 ± 1.97) (Supplementary Figures 4C–D; 5C–D; Supplementary Tables S12a, d). ANOVA detected significant differences among genotypes (*AtOFP2*: F = 113.4; *AtOFP17*: F = 106.5; Fcrit = 3.22; P < 0.001; Supplementary Table S12b, d). Tukey’s test confirmed that both knock-down and over-expression lines differed strongly from WT (q ≈ 18 for both genes), while *amiRNA* vs. *Oex* comparisons were non-significant (q < 1; Supplementary Table S12c, f). R² values (∼0.89 for both) indicate genotype as a major contributor to stem trichome density, though additional factors may also influence this trait.

### Plant height at maturity

Height of mature plants was measured at seed set (∼10 weeks for *amiR-ofp2* and WT; ∼15 weeks for *oex-ofp2*). *amiR-ofp2* plants were significantly taller (31.5 ± 2.55 cm) than both WT (21.4 ± 2.72 cm) and *oex-ofp2* (21.7 ± 2.25 cm; Figure 3H, I; Supplementary Table S13a). *amiR-ofp17* plants also showed increased height (25.93 ± 2.91 cm) compared to WT (22.13 ± 2.23 cm) and *oex-ofp17* (15.93 ± 1.48 cm), though the genotype effect was weaker (Figure 4G, J; Supplementary Table S13d). ANOVA revealed significant differences (*AtOFP2*: F = 78.41; *AtOFP17*: F = 73.10; Fcrit = 3.22; P < 0.001; Supplementary Table S13b, e). Tukey’s test for *AtOFP2* showed amiRNA lines differed strongly from both WT and oex (q = 15.59 and 15.07), while WT vs. Oex was non-significant (q = 0.51; Supplementary Table S13f). For *AtOFP17*, all pairwise comparisons were significant (q = 6.44–16.94), exceeding q₀.₀₅crit = 3.44 and q₀.₀₁crit = 4.35 (Supplementary Table S13f). R² values were moderate (*AtOFP2*: 0.79; *AtOFP2*: 0.78), indicating other factors besides genotype also influence plant height.

### Number of primary and secondary branches

The trait values were counted in plants at the time of maturity (n=15). In *amiR-ofp2*, the number of primary branches (4.33 ± 0.81) was higher than WT (3.73 ± 0.79), while *oex-ofp2* had fewer branches (2.73 ± 0.73; Figure 3H, J; Supplementary Table S14a). A similar pattern was observed for *amiR-ofp17* (4.33 ± 0.89), WT (4 ± 1.07), and *oex-ofp17* (2.73 ± 0.88; Figure 4G, I; Supplementary Table S14f). ANOVA indicated modest but significant differences (*AtOFP2*: F = 16.33; *AtOFP17*: F = 11.73; Fcrit = 3.22; P < 0.001; Supplementary Table S14b, g). Tukey’s test showed oex lines differed significantly from both amiRNA and WT (*AtOFP2*: q = 8.00 and 5.00; *AtOFP17*: q = 6.49 and 5.14; q₀.₀₅crit = 3.44; q₀.₀₁crit = 4.35). amiRNA vs. WT comparisons were non-significant in both genes (q < 3.1; Supplementary Table S14c, h). R² was low (*AtOFP2*: 0.44; *AtOFP17*: 0.35), suggesting that genotype alone explains little of the variation in branch number.

Number of secondary branches was counted at 15.13 ± 2.23 in *amiR-ofp2*, 11 ± 1.69 in WT, and 11.8 ± 2.336 in *oex-ofp2* (Figure 3H, J; Supplementary Table S14a). Corresponding values were 12.46 ± 2.29 (*amiR-ofp17*), 12.13 ± 2.445 (WT) and 7.266 ± 1.83 (*oex-ofp17*; Figure 4G, I; Supplementary Table S14f). ANOVA detected modest but significant differences (*AtOFP2*: F = 16.27; *AtOFP17*: F = 26.11; Fcrit = 3.22; P < 0.001; Supplementary Tables S14d, i). Tukey’s test for *AtOFP2* showed amiRNA differed significantly from both WT and oex (q = 7.60 and 6.13), while WT vs. oex was non-significant (q = 1.47). In *AtOFP17*, oex plants differed significantly from both amiRNA and WT (q = 8.54–9.13), while amiRNA vs. WT was non-significant (q = 0.59; Supplementary Table S14e, j). R² values were low (*AtOFP2*: 0.44; *AtOFP17*: 0.55), suggesting minimal genotype-phenotype correlation for secondary branch number. Taken together with primary branch counts, these results indicate that branching patterns are only weakly influenced by genotype compared to other shoot traits such as height or rosette area.

### Seed related traits

#### Silique length and number of seeds per silique

Length was measured in 15 randomly selected siliques at maturity from the tip of the beak to the base of attachment with the pedicel. Seeds per siliques were also measured in the same siliques. amiRNA lines produced both the longest siliques and the highest seed counts in *AtOFP2* and *AtOFP17*. Silique length averaged 14.02 ± 0.80 mm with 50.66 ± 3.06 seeds/silique in *amiR-ofp2*, compared to 9.44 ± 0.72 mm and 39.13 ± 1.84 seeds/silique in WT, and 9.66 ± 0.62 mm silique length with 39.26 ± 1.83 seeds/silique in *oex-ofp2* (Figure 5E, J and K; Supplementary Table S15a). Silique length and seeds/silique values were 13.42 ± 0.99 mm with 47.06 ± 2.57 seeds/silique in *amiR-ofp17*, 9.83 ± 0.70 mm with 40.6 ± 2.66 seeds/silique in WT, and 8.78 ± 0.90 mm with 26.13 ± 2.47 seeds/silique in *oex-ofp17* (Figure 6E, H and I; Supplementary Table S15f).

**Figure 5:**
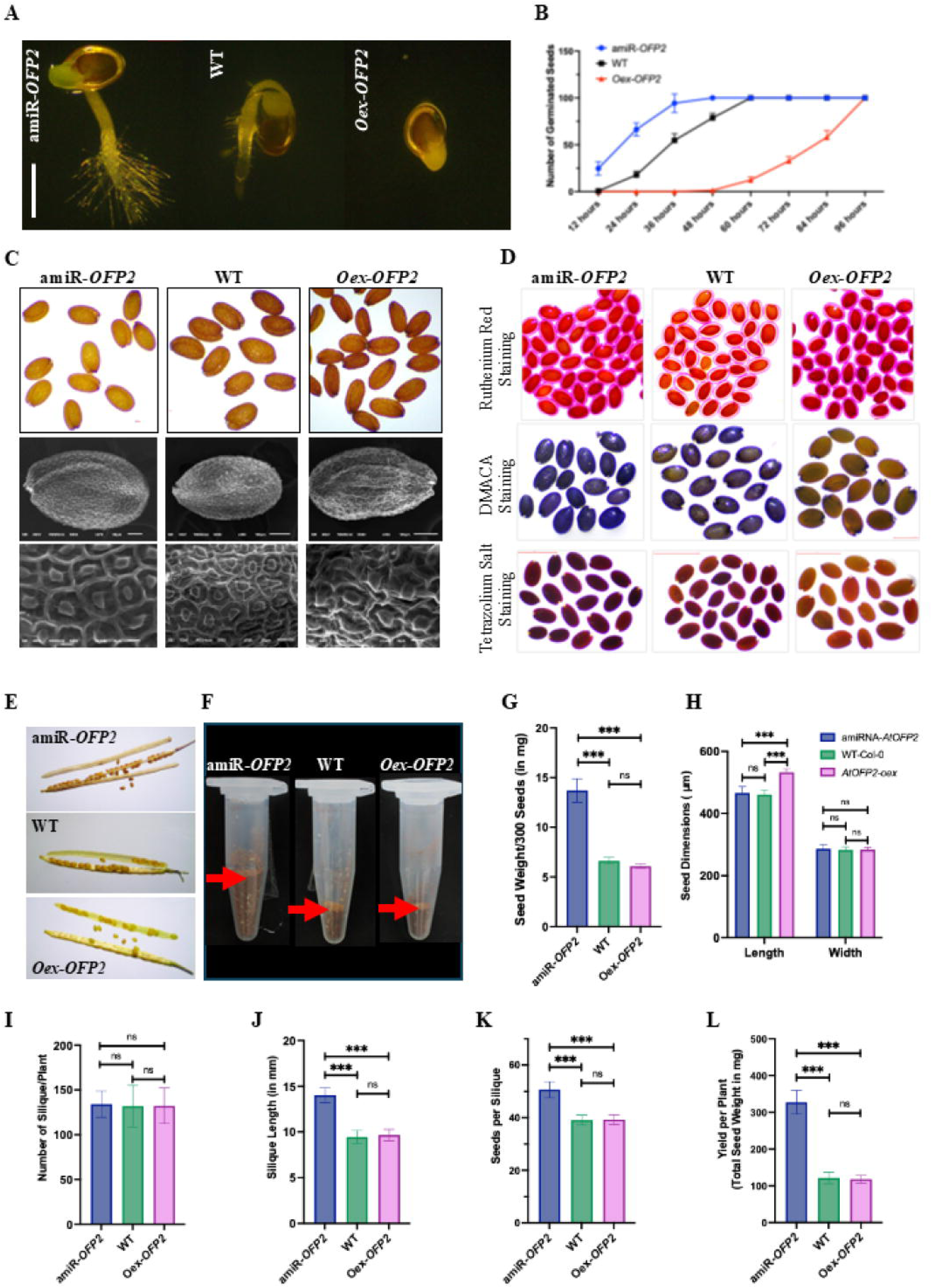
Comparison of physical and physiological traits of seeds and final yield between WT, knock-down (*amiR-OFP2*) and over-expression (*oex-OFP2*) plants. **A)** Germinated seeds after 48-hours of plating. **B)** Percent germination of *amiR-ofp2*, WT and *oex*- *OFP2* seeds. **C)** comparison of seed characteristics between WT, knock-down (*amiR-OFP2*) and over-expression (*oex-OFP2*): Top panel: Dry seeds. Middle panel: Scanning electron micrograph of whole seeds. Bottom panel: Scanning electron micrograph showing close-up of seed coat epidermal cells. **D)** Staining of seeds - Top panel: Mucilage of hydrated seeds stained with ruthenium red. Middle panel: DMACA staining for Proanthocyanidins in seed coat, Bottom panel: Tetrazolium stain for permeability of seed coat. **E)** Siliques from mature plants of *amiR-ofp2*, WT and *oex-ofp2* to show silique length and seeds per silique. **F)** representation of total yield in the form of seeds obtained from a single plant. **G, H, I, J, K, L)** Graphical representation of statistical comparisons of weight of 300 dry seeds; dimensions of seeds; number of siliques per plant, silique length, seeds per silique and yield per plant. Bars in all graphs represent mean ± SD, ns = no significant difference, *p=0.033, **p=0.02, ***p<0.001.

**Figure 6:**
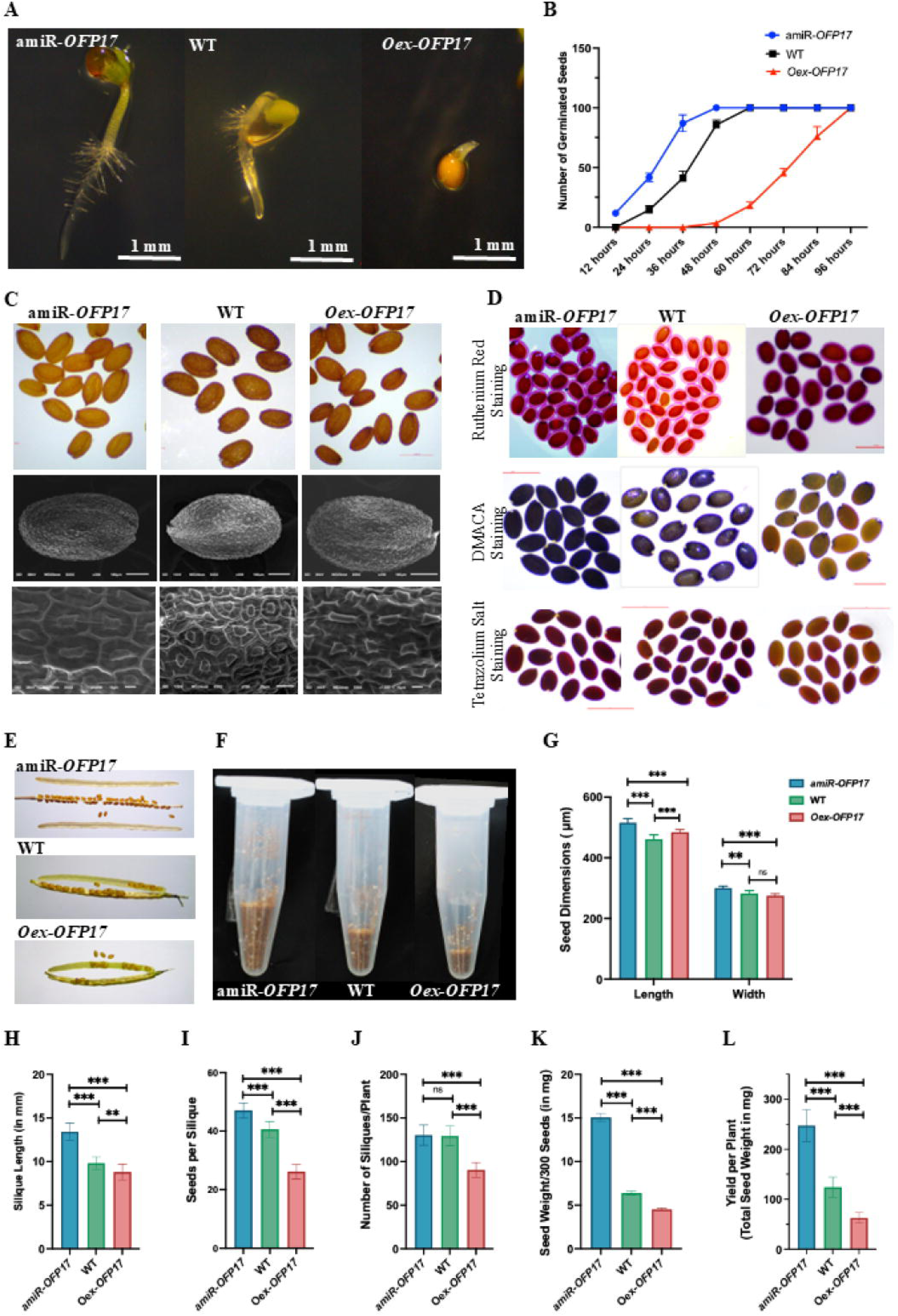
Comparison of physical and physiological traits of seeds and final yield between WT, knock-down (*amiR-OFP17*) and over-expression (*oex-OFP17*) plants. **A)** Germinated seeds after 48-hours of plating. **B)** Percent germination of *amiR-ofp17*, WT and *oex-ofp17* seeds. **C)** comparison of seed characteristics between WT, knock-down (*amiR-OFP17*) and over-expression (*oex-OFP17*):Top panel: Dry seeds. Middle panel: Scanning electron micrograph of whole seeds. Bottom panel: Scanning electron micrograph showing close-up of seed coat epidermal cells. **D)** Staining of seeds - Top panel: Mucilage of hydrated seeds stained with ruthenium red. Middle panel: DMACA staining for Proanthocyanidins in seed coat, Bottom panel: Tetrazolium stain for permeability of seed coat. **E)** Siliques from mature plants of *amiR-ofp17*, WT and *oex-ofp17* to show silique length and seeds per silique. **F**) representation of total yield in the form of seeds obtained from a single plant. **G, H, I, J, K, L)** Graphical representation of statistical comparisons of dimensions of seeds, silique length, seeds per silique, number of siliques per plant, weight of 300 dry seeds and yield per plant. Bars in all graphs represent mean ± SD, N=10, ns = no significant difference, *p=0.033, **p=0.02, ***p<0.001.

ANOVA confirmed strong genotype effects for both traits (*AtOFP2*: F = 193.2 for length, 122.2 for seeds; *AtOFP17*: F = 115 for length, 260.1 for seeds; Fcrit = 3.22; P < 0.001; Supplementary Tables S15b, S15d, S15g, S15i). Tukey’s test showed significant separation of amiRNA from WT and oex in all cases (*AtOFP2*: q = 19.03–24.65; *AtOFP17*: q = 9.73–31.5), while WT vs. oex was non-significant for *AtOFP2* silique length and seed number (q ≈ 1; Table Supplementary S15c, S15e, S15h, S15j). R² values were high for silique length (*AtOFP2*: 0.90; *AtOFP17*: 0.85) and for seed number (*AtOFP2*: 0.85; *AtOFP17*: 0.92), indicating genotype is a major determinant of these traits. Both genes showed a consistent amiRNA > WT > oex pattern, linking *OFP2/17* knockdown to enhanced silique development and reproductive output.

### Number of siliques and seed yield per plant

Silique counts at harvest (n = 15 plants) were largely similar among *AtOFP2* genotypes: *amiR-ofp2* (134.06 ± 14.76), WT (131.86 ± 23.05), and *oex-ofp2* (132.2 ± 19.89; Figure 5I; Supplementary Table S16a). In *AtOFP17* genotypes, count in *amiR-ofp17* (130.26 ± 11.65) and WT (129.4 ± 11.09) were comparable, while *oex-ofp17* had reduced count (90.33 ± 8.06) (Figures 6J; Supplementary Tables S16f). ANOVA found no significant genotype effect in *AtOFP2* (F = 0.055; Fcrit = 3.22; P = 0.95; Supplementary Table S16b) with all Tukey’s q-values < 0.5 (Supplementary Table S16c), and R² near zero (0.0026), indicating silique number is independent of genotype in *AtOFP2* (Supplementary Table S16c). In contrast, *AtOFP17* showed significant differences (F = 70.31; P < 0.001), with *oex-ofp17* differing from both amiRNA and WT (q ≈ 14.4–14.7), while amiRNA vs. WT remained non-significant (q = 0.32) (Supplementary Tables S16g-h).

Seed yield per plant followed a clear genotype-dependent trend. Highest seed yield (327.26 ± 32.06 mg) was observed in *amiR-ofp2* whereas the yield in WT (121.2 ± 15.70 mg) and *oex-ofp2* (117.86 ± 11.03 mg) were much lower and nearly identical (Figure 5F; L; Supplementary Table S16a). Similarly, seed yield was highest (247.33 ± 32.06 mg) in *amiR-ofp17*, followed by WT (124.2 ± 20.14 mg) and *oex-ofp17* (62.8 ± 10.81 mg) (Figures 6F, L; Supplementary Tables S16f).

ANOVA confirmed strong genotype effects for both genes (*AtOFP2*: F = 452.1; *AtOFP17*: F = 256.3; Fcrit = 3.22; P < 0.001; Supplementary Table S16d, S16i). Tukey’s test showed amiRNA significantly differed from both WT and oex in each gene (*AtOFP2*: q = 36.53–37.12; *AtOFP17*: q = 20.98–31.44), while WT vs. oex was non-significant in *AtOFP2* (q = 0.59) but significant in *AtOFP17* (q = 10.46; Supplementary Table S16e, S16j).

### Seed morphology and weight

Dry mature seeds of WT and mutants were observed under stereo-microscope and imaged. The seeds of *amiR-ofp2* and *amiR-ofp17* seeds were light brown as compared to WT, whereas those of *oex-ofp2* and *oex-ofp17* seeds were darker (upper panels in Figures 5C, 6C). The weight of 300 seeds was highest in *amiR-ofp2* (13.7 ± 1.8 mg) compared to WT (6.62 ± 0.36 mg) and *oex-ofp2* (6.08 ± 0.24 mg; Figure 5G; Supplementary Table S17a). The corresponding data on seed weight were 15.06 ± 0.45 mg in *amiR-ofp17*, which was significantly heavier than WT (6.36 ± 0.215 mg) and *oex-ofp17* (4.51 ± 0.15 mg; Figure 6K; Supplementary Table S17d). One-way ANOVA confirmed highly significant variation across genotypes (*AtOFP2*: F = 102.8; *AtOFP17*: F = 1051; Fcrit = 3.22; P < 0.001; Supplementary Tables S17b, S17e). In *AtOFP2*, Tukey’s test identified significant differences between amiRNA and both control and over-expression (q = 16.88–18.17), while WT and oex did not differ significantly (q = 1.29). In *AtOFP17*, all pairwise comparisons were significant, with exceptionally high q-values for amiRNA vs. oex (q = 60.71) and WT vs. oex (q = 10.66) (Supplementary Table S17c, S17f).

Strong goodness-of-fit (*AtOFP2*: R² = 0.97; *AtOFP17*: R² = 0.99) underscores genotype as a dominant factor influencing seed weight. Overall, both genes displayed a consistent trend, seed mass was elevated in knockdown lines, intermediate in WT, and lowest in overexpression lines.

### Seed dimension

To assess seed morphology, dry mature seeds were first examined under a stereo-microscope and then imaged using scanning electron microscopy for precise measurement (Figures 5C, 6C, lower panels). ImageJ analysis revealed notable differences in seed length among genotypes. In the *AtOFP2* genotypes, *oex-ofp2* seeds were longest (532.4 ± 11.5 μm), followed by *amiR-ofp2* (465.7 ± 20.8 μm) and WT (461.1 ± 14.2 μm). Seed width, however, remained largely unchanged across genotypes, averaging 286.4 ± 13.0 μm in *amiR-ofp2*, 283.5 ± 7.4 μm in *oex-ofp2*, and 282.6 ± 9.5 μm in WT (Figure 5H; Supplementary Table S18a). Statistical analysis via one-way ANOVA confirmed significant variation in seed length (F = 46.63, P < 0.001, F_crit = 3.354; Supplementary Table S18b), with Tukey’s test identifying all pairwise differences as significant: *amiR-ofp2* vs. WT (q = 13.6), *amiR-ofp2* vs. *oex-ofp2* (q = 7.86), and WT vs. *oex-ofp2* (q = 5.73), all above the q_0.01 critical threshold of 4.25 (Supplementary Table S18c). For width, ANOVA also found significant differences (F = 28.37, P < 0.001), but Tukey’s test indicated only the *amiR-ofp2* vs. *oex-ofp2* (q = 10.37) and *amiR-ofp2* vs. WT (q = 7.3) comparisons were significant, while WT vs. *oex-ofp2* (q = 3.069) fell below the q_0.01 cutoff (Supplementary Tables S18d-e).

In contrast, the *AtOFP17* lines displayed a reversed pattern in seed length. *amiR-ofp17* seeds were longest (515.4 ± 13.8 μm), followed by *oex-ofp17* (484.0 ± 9.3 μm), with WT being shortest (461.1 ± 14.2 μm). For seed width, measurements were variable: 300.2 ± 6.54 μm (*amiR-ofp17*), 282.6 ± 9.5 μm (WT), and 275.2 ± 6.54 μm (*oex-ofp17*; Figure 6G; Supplementary Table S18f). One-way ANOVA for seed length showed strong significance (F = 62.35, P < 0.001), and Tukey’s q-values confirmed significant differences between *amiR-ofp17* vs. WT (q = 13.2) and *amiR-ofp17* vs. *oex-ofp17* (q = 14.11), while WT vs. *oex-ofp17* (q = 0.91) was not significant (Supplementary Tables S18g-h). For width, however, ANOVA yielded a non-significant result (F = 0.3766, P = 0.690), and all pairwise Tukey’s q-values (1.174, 0.896, and 0.278) were well below the critical threshold (Supplementary Tables S18i-j).

Goodness-of-fit values reinforced these findings: in the *AtOFP2* lines, R² values were 0.822 (length) and 0.0271 (width), suggesting genotype strongly influences seed length but not width. In *AtOFP17*, seed length also showed high genotype dependence (R² = 0.87), while seed width remained genotype-independent (R² = 0.016). These results collectively suggest that *AtOFP*2 and *AtOFP17* have distinct regulatory roles in controlling seed elongation, but exert limited influence on seed width.

### Life span

We compared the time taken to complete the life cycle of WT and mutant plants at T3 generation (n=5) under similar growth conditions. The plants of *amiR-ofp2* completed their life cycle in approximately 10 ± 0.70 weeks, while WT took 11.2 ± 0.836 weeks, and *oex-ofp2* required the longest time of 14.8 ± 0.836 weeks (Figure 7K; Supplementary Table S19a). In *AtOPF17*, average life span was 10.4 ± 0.55 weeks (*amiR-ofp17*) to 11.1 ± 0.74 (WT) and 11.4 ± 0.55 weeks (*oex-ofp17*), (Supplementary Table S19d). Genotype has a significant effect on life span, particularly in *AtOFP2* lines, but not as pronounced in *AtOFP17*. One-way ANOVA showed a significant effect of genotype (F = 49.26 > Fcrit = 3.885, *P* < 0.001). Tukey’s test indicated significant differences between *oex-ofp2* and both WT (q = 10.12) and *amiR-ofp2* (q = 13.49), but not between *amiR-ofp2* and WT (q = 3.372) (Supplementary Table S19b-c).

**Figure 7:**
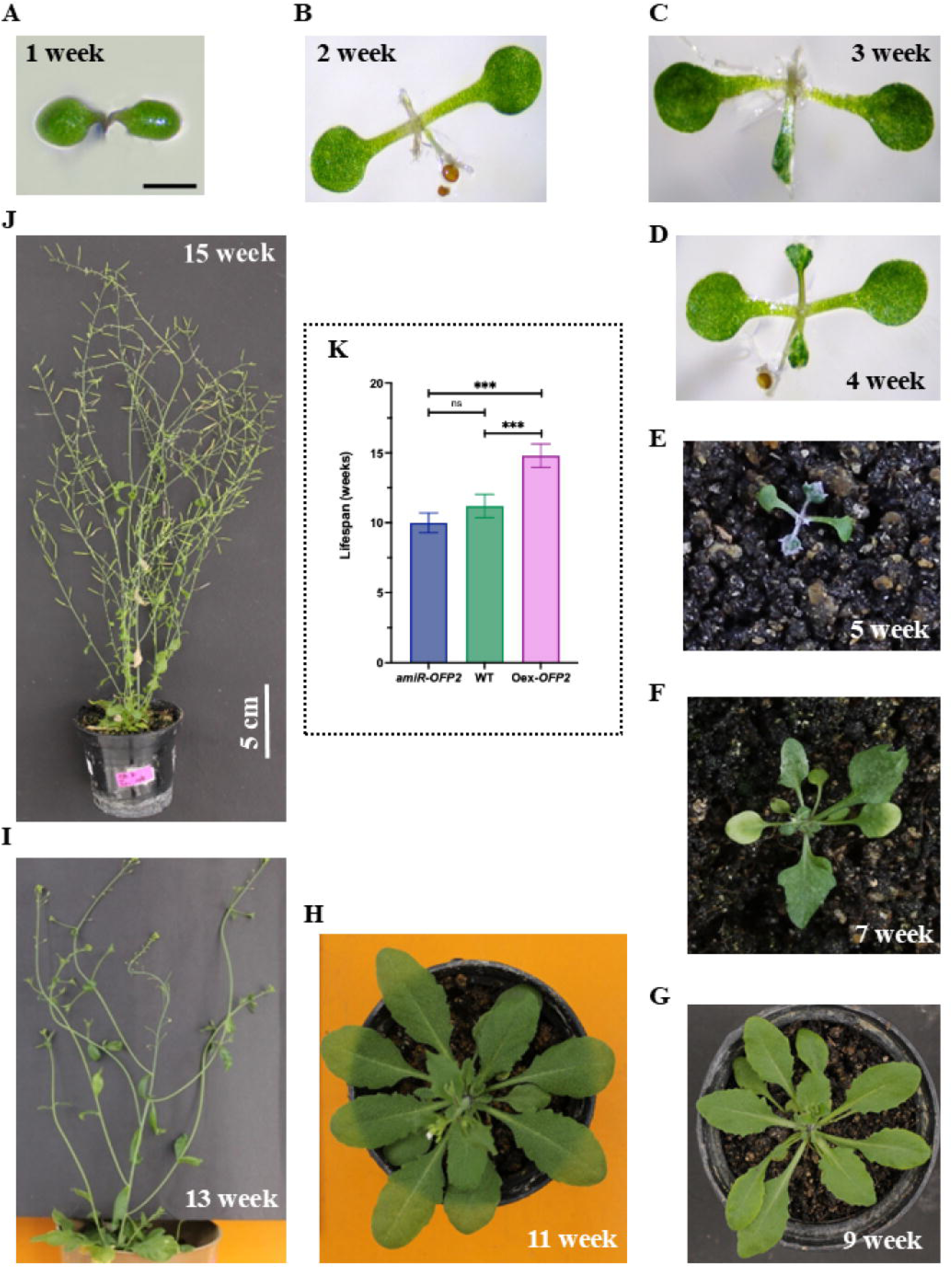
[A-J]. Stages in life-cycle of *AtOFP2* over-expression plant from week-1 to week-15. **[K]** Graphical representation of statistical comparisons of life spans of *amiR-ofp2*, WT and *oex-ofp2* plants. Bars represent mean ± SD, N=5, ns = no significant difference, *p=0.033, **p=0.02, ***p<0.001

In *AtOFP17* lines, life span differences were subtler: 10.4 ± 0.55 weeks (*amiR-ofp17*), 11.1 ± 0.74 weeks (WT), and 11.4 ± 0.55 weeks (*oex-ofp17*). ANOVA showed a marginally significant group difference (F = 4.739 > Fcrit = 3.885). Tukey’s test found a slightly significant difference only between *amiR-ofp17* and *oex-ofp17* (q = 4.334), while other comparisons were not significant (WT vs. *amiR-ofp17*: q = 2.528; WT vs. *oex-ofp17*: q = 1.806) (Supplementary Tables S19e-f). Thus, genotype strongly influenced lifespan in *AtOFP2* lines, but only mildly in *AtOFP17* lines.

### Dormancy

Seed dormancy was assessed by tracking the percent germination of non-stratified seeds in WT and mutant lines. One hundred seeds of each genotype were placed on ½ strength MS agar plates and exposed to a controlled light-dark cycle (2h light, 8h dark on day-one, then 16h light/8h dark from day-two onwards). Seeds were scored for radicle emergence at 12-hour intervals, with full germination occurring within 96 hours; germinated seeds were photographed after every 48 hours for comparison. In *amiR-ofp2*, seed germination started within 12 hours (av. 24.67 ± 7.06%) and reached 100% in 48 hours. In WT, germination rate at 12 hours was only 0.67% which reached 17.67 ± 7.09 in 24 hours and 100% in 60 hours. In *oex-ofp2*, germination started only after 36 hours and average germination after 48 hours was only 1.33 ± 1.53%, with 100% germination only by 96 hours. The knockdown mutants of *AtOFP2* showed reduced dormancy compared to WT, while overexpression lines displayed increased dormancy (Figure 5 A, B; Supplementary Tables S20a-b).

Similarly, in *amiR-ofp17*, germination commenced within 12 hours (11.67 + 2.08%) and reached 100% within 48 hours. In WT, germination started after 12 hours (14.33 ± 3.51%) reaching 100% after 60 hours. In *oex-ofp17*, seed germination started only after 36 hours and reached 3.33 ± 1.52% after 48 hours; 100% seed germination was observed at 96 hours (Figure 6A, B; Supplementary Table S20c-d).

### Seed shape and epidermal cell structure

Scanning electron microscopy (SEM) revealed morphological differences between the seeds of different genotypes. *amiR-ofp2* seeds were rounder and larger than WT, while *oex-ofp2* seeds exhibited an elongated and shrunken morphology. The epidermal cell structure of the seedcoat also showed differences, with *amiR-ofp2* seeds having more compact cells and thicker radial walls than WT, whereas *oex-ofp2* seeds displayed disrupted cell walls, enlarged columella, and disoriented cellulosic microfibrils. Seed coat mucilage cells or epidermal cells of seed wall of wild type *A. thaliana* seeds are hexagonal cells with a “volcano shaped” columella in centre. Due to secondary cell wall synthesis, walls of cells are distinctly visible in the scanning electron micrograph. These cells contain mucilage between plasma membrane and cell wall, and form mucilage pockets surrounding columella. Cellulosic microfibrils radiating from the columella appear as fine lines. These microfibrils are responsible for cell shape. Cortical microtubules ensure the correct arrangement of cellulosic microfibrils (Windsor et al, 2000). In *amiR-ofp2* seeds, radial walls of epidermal cells were more prominent and thickened than WT seeds exhibiting more compactness in the cells. On the contrary, epidermal cells in *oex-ofp2* seeds had disrupted cell walls, enlarged columella and disoriented cellulosic microfibrils (Figure 5C-lower panels).

SEM images of *amiR-ofp17*, WT, and *oex-ofp17* seeds revealed similar trends. *amiR-ofp17* seeds had compact epidermal cells with prominent radial walls, while *oex-ofp17* seeds exhibited disrupted cell walls, enlarged columella, and elongated, rectangular cells. In *oex-ofp17,* the radial walls were found to be disrupted, but the impact was lesser than that observed in *oex-ofp2* (Figure 6C-lower panels).

### Mucilage release from testa

The mucilage release properties of seeds from *AtOFP2* and *AtOFP17* reverse genetic lines were examined to determine how secondary cell wall modifications affect mucilage extrusion. *A. thaliana* seeds are known to release non-adherent mucilage immediately upon contact with moisture, while the adherent mucilage takes time to form a halo around the seeds (Western et al., 2000, Arsovski et al, 2010). Seeds from *amiR-ofp2*, *oex-ofp2*, and WT were soaked in water for 1 hour and stained with ruthenium red, which binds to acidic polysaccharides, producing a pink coloration. Seeds from all constructs released ample mucilage and formed halos, with no observable differences between the lines (upper panels in Figure 5D).

Seeds from *amiR-ofp17*, *oex-ofp17*, and WT were similarly soaked for 1 hour and stained. While *amiR-ofp17* and WT seeds showed complete mucilage release, indicated by the formation of halos around the seeds, *oex-ofp17* seeds displayed spikes radiating from the seed surface. This observation suggests that mucilage was still being extruded from the seed wall after hydration, indicating that mucilage release was slower in *oex-ofp17* seeds compared to *amiR-ofp17* and WT seeds (figure 6D, upper panel). These results highlight the altered dynamics of mucilage release in *oex-ofp17*, possibly due to changes in the seed coat structure.

### Proanthocyanidin (PA) staining

Proanthocyanidin (PA) is a polymeric flavanol and its precursor accumulates in the endothelial cells of seed coat. After harvesting, oxidation of PA precursor imparts dark brown colour to the seeds. As we had observed colour difference in the seeds of knock-down and over-expression lines, we stained the dried seeds with DMACA (p-dimethylaminocinnamaldehyde) reagent which binds to flavanols and gives a purple to blue colour. Intensity of colour indicates the presence of proanthocyanidins in endothelial cells of seed coat. Proanthocyanidin content was analyzed using DMACA staining, which revealed a gradient in PA levels across the genotypes. *amiR-ofp2* seeds showed the highest intensity of purple-blue staining, indicating maximum PA accumulation, followed by WT seeds with moderate staining. *oex-ofp2* seeds exhibited minimal staining, localized primarily at the chalazal end. A similar trend was observed for *amiR-ofp17* and *oex-ofp17* lines (middle panels in Figure 5D, 6D).

### Seed coat permeability

Permeability of seed coat is an essential prerequisite for seed germination. Lesser permeability of the seed coat to water can hamper germination. To observe the water uptake properties of seeds, we dipped the seeds in tetrazolium chloride salt for 1 day. TZ (2,3,5 triphenyl tetrazolium chloride) is a substance which can penetrate both living and dead cells, but only respiring cells can convert TZ to a carmine red coloured water-insoluble formazan. Intensity of colour depicts the amount of TZ absorbed and converted into formazan. Seed coat permeability was assessed using tetrazolium chloride (TZ) uptake. The intensity of carmine red formazan formation indicated that *amiR-ofp2* seeds had the highest permeability, followed by WT, while *oex-ofp2* seeds had the least permeability. Similarly, *amiR-ofp17* seeds exhibited greater permeability than WT, and *oex-ofp17* seeds absorbed the least TZ, reflecting reduced seed coat permeability in overexpression lines (Figure 5D, 6D lower panel). These findings collectively demonstrate that altering *AtOFP2* and *AtOFP17* expression impacts seed dormancy, morphology, mucilage release, PA content, and seed coat permeability. Knockdown mutants tend to reduce dormancy and enhance seed coat permeability, while overexpression lines show opposite effects (lower panels in Figure 5D, 6D).

### *AtOFP2* has a stronger repressive effect than *AtOFP17*

Phenotypic analysis of *Arabidopsis thaliana* plants overexpressing *AtOFP2* (*oex-ofp2*) and *AtOFP17* (*oex-ofp17*) unequivocally demonstrates that both genes broadly exert repressive effects on various developmental traits (Figure 10). However, a critical aspect of this analysis is the revelation of distinct and nuanced differences in the degree and specific nature of these repressive effects, indicating unique functional specificities or underlying regulatory mechanisms for each gene. These distinctions are crucial for understanding the precise roles of these paralogous genes in plant development. During the juvenile stage of development, overexpression of both *AtOFP2* and *AtOFP17* resulted in a significant repression of seedling growth. However, a quantitative assessment revealed that growth stunting was markedly more pronounced in *oex-ofp2* lines compared to *oex-ofp17* lines. This disparity is clearly illustrated by specific quantitative data: the average length of roots in *oex-ofp17* was measured at 0.93 cm±0.284, whereas in *oex-ofp2*, the average root length was substantially shorter at only 0.41 cm±0.091. This quantitative difference signifies that the roots of *oex-ofp17* seedlings were approximately 2.2-fold longer than those of *oex-ofp2*, despite both being significantly stunted when compared to wild-type plants. This compelling quantitative evidence strongly supports the notion of a differential repressive strength. Across all seedling traits evaluated, with the sole exception of hypocotyl length, *AtOFP2* overexpression consistently resulted in more severe phenotypic effects than *AtOFP17* overexpression. This suggests a broad and pervasive repressive impact of *AtOFP2* on overall seedling vigor and the fundamental processes of early developmental establishment. Beyond root growth, seed germination was quantitatively more delayed in *oex-ofp2* lines compared to *oex-ofp17* lines further reinforcing stronger and more comprehensive repressive impact of *AtOFP2* at the very initial stages of plant development. Reversal of repression of both genes in knock-down lines caused higher seed yield compared to WT and among both knock-down mutants seed yield was considerably higher in *amir-ofp2* than *amir-ofp17* mutants. Most importantly, ∼90% seedling of *oex-ofp2* mutant background died at juvenile stage when transferred to pots and only ∼10% seedling could make it to a mature plant. No seedling death was observed for *oex-ofp17* mutants. The provision of such precise quantitative data, like the average root lengths and the explicit statement about the widespread impact on seedling traits, higher seed yield in knock-down lines and seedling death in over-expression mutants offers compelling and measurable evidence for *AtOFP2* being a significantly stronger repressor at the seedling stage.

### Strong repression at juvenile stage caused heterochronic shift in life cycle

A particularly striking and critical observation in *oex-ofp2* mutants was the markedly reduced survival rate of seedlings (<10%) upon transfer from agar plates to soil, indicating a severe developmental bottleneck or cellular stress response induced by sustained *AtOFP2* overexpression leading to high mortality. On the agar plates, all other genotypes grew well and could be transferred to soil after 2 weeks of germination. *oex-ofp2* mutants were barely transferable to soil due to extremely small roots even after 3^rd^ week and took longer than usual time for root establishment in soil. These seedlings exhibited a prolonged dormant phase, characterized by remaining at the four-leaf stage for nearly one month. During this extended period, newly emerging pairs of leaves consistently underwent degeneration (Figure 3G). Remarkably, seedlings that managed to overcome this dormancy eventually resumed normal growth and development, exhibiting a pattern strikingly similar to that of wild-type plants. This initial inhibition of growth at juvenile stage shifts the time of growth and development causing a heterochronic shift and expanded the life span (eg. 11.2 ± 0.8 week for wild type Col-0 v/s 14.8 ± 0.8 for *oex-ofp2*; Figure 7). Wild type attains maturity in 7-8 weeks while *oex-ofp2* plants attain maturity in 11 - 12 weeks. WT plants complete their life cycle in 10 -12 weeks whereas *oex-ofp2* plants takes 15 – 16 weeks (Figure 7).

### Transcriptome data analysis results

A comparative analysis of reverse genetic mutants revealed phenotypes of *ofp2* stronger than those of *ofp17*. Among all the phenotypic classes, seedling root system architecture was one of the most significantly affected. We therefore analyzed the underlying changes in gene expression pattern in roots of 7-day old seedlings of WT, *oex-ofp2* and *amiR-ofp2* via transcriptome profiling. Total RNA was extracted and subjected to paired-end short-read based RNA sequencing using Illumina NovaSeq 6000 platform. Differential gene expression analysis was carried out using DESeq2 in R, and significantly deregulated genes were visualized using volcano plots (Figure 8A and 8B).

**Figure 8:**
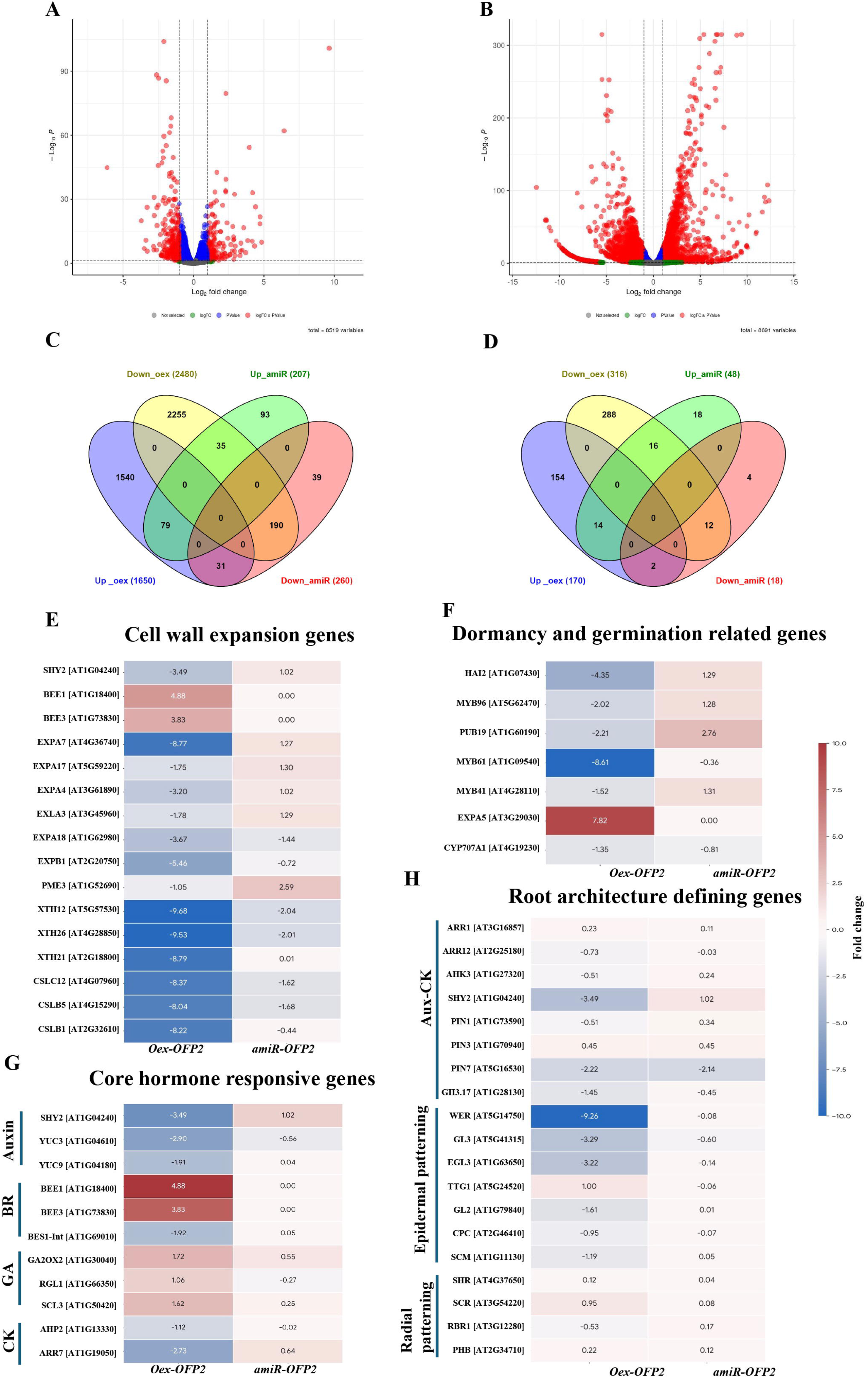
RNA-seq analysis of roots of 7-day old seedlings from WT, *oex-OFP2* and *amiR-OFP2* lines. **A)** Volcano plot showing differentially expressed genes (DEGs) from WT vs *amiR-OFP2*. **B)** Volcano plot showing differentially expressed genes (DEGs) from WT vs *oex-OFP2*. **C)** Venn diagram showing common and unique significantly deregulated genes among *amiR-OFP2* and *oex-OFP2* genotypes. **D)** Venn diagram showing common and unique significantly deregulated transcription factors in *amiR-OFP2* and *oex-OFP2* genotype. **E, F, G, H)** – Heat map of important gene clusters deregulated in *oex-ofp2* and *amiR-ofp2* mutants representing (i) cell wall expansion gene, (ii) dormancy and germination related genes, (iii) core hormone responsive genes (iv) root architecture defining genes.

In *oex-ofp2*, a total of 8,691 genes were differentially expressed relative to WT. Genes exhibiting a log₂ fold change (log₂FC) > 1 or < –1 were selected for downstream analysis (Supplementary Table S21a). Among these, 1,650 genes were significantly up-regulated, while 2,480 genes were significantly down-regulated (Supplementary Table S21b, c). In *amiR-ofp2* seedlings, 8,519 genes were deregulated compared to WT. Of these, 208 genes were significantly up-regulated and 260 genes were significantly down-regulated (Supplementary Table S22a–c), indicating a more restrained transcriptional response in the knockdown background. Comparative analysis between the two mutant backgrounds revealed that 31 genes were up-regulated in *oex-ofp2* and down-regulated in *amiR-ofp2*, while 35 genes displayed the opposite trend, being down-regulated in *oex-OFP2* and up-regulated in *amiR-OFP2* (Figure 8C). In addition, 1,540 genes were uniquely up-regulated and 2,255 genes uniquely down-regulated in *oex-OFP2*, whereas 93 and 39 genes were uniquely up- and down-regulated, respectively, in *amiR-OFP2* (Supplementary Table 22b–e).

Given that *OFP2* functions as a transcription factor, we specifically examined transcription factor (TF) genes within the significantly deregulated gene sets. In *oex-OFP2*, 170 TFs were significantly up-regulated and 316 TFs were significantly down-regulated (Supplementary Table S21). In contrast, *amiR-OFP2* plants showed 48 up-regulated and 18 down-regulated TFs (Supplementary Table S22). Analysis of common and unique TFs among both genotypes found two TFs were up-regulated in *oex-OFP2* and down-regulated in *amiR-OFP2*, whereas 16 TFs were down-regulated in *oex-OFP2* and up-regulated in *amiR-OFP2* (Figure 8D; Supplementary Table S21-22). Furthermore, 154 TFs were uniquely up-regulated and 280 TFs uniquely down-regulated in *oex-OFP2*, while 18 and 4 TFs were uniquely up- and down-regulated, respectively, in *amiR-OFP2* (Supplementary Table S21-22).

We used the AGRIS AtTFDB to classify the deregulated TFs from both *oex-ofp2* and *amiR-ofp2* lines. A total of 40 TF families are deregulated out of which 27 TF families are deregulated in both *oex-ofp2* and *amiR-ofp2* lines. These include *ABI3VP1, AP2-EREBP, ARF, ARR-B, BBR/BPC, bHLH, bZIP, BZR, C2C2-CO-like, C2C2-Dof, C2C2-GATA, C2H2, CCAAT-HAP2, G2-like, GRAS, GRF, Homeobox, HSF, MADS, MYB, NAC, RAV, SBP*, *TCP, Trihelix, WRKY* and *ZF-HD*. The *C2C2-YABBY* family occurs exclusively in upregulated TF category of *oex-ofp2*. The *Orphan* TF family was common between downregulated category of *amiR-ofp2* and upregulated category of *oex-ofp2*. Two families, *VOZ-9* and *Whirly* are shared between upregulated categories of TF of *amiR-ofp2* and *oex-ofp2.* The *NLP* family was common between downregulated categories of *amiR-ofp2* and *oex-ofp2*. Five TF families (*AtRKD, CAMTA, CCAAT-DR1, HRT, REM*) were shared between upregulated category of *amiR-ofp2* and downregulated category of *oex-ofp2*. The *CPP, E2F-DP*, and *EIL3* family were shared between upregulated category of *amiR-ofp2*, and downregulated categories of both *amiR-ofp2* and *oex-ofp2* (Supplementary table S21 and S22). Gene ontology and KEGG pathway analysis using significantly deregulated genes revealed alteration mainly in plant hormone signal transduction pathway (Supplementary figure 6) and circadian rhythm pathway (supplementary figure 7).

The read count of *OFP2* is 110.33 ± 5.5 in wild type Col-0; in comparison, the maximum read count in wild type is 220150.66 and average read count is 881.83. This indicates that *OFP2* in its native form is a relatively less expressed and even this low level is enough to exert its effect. In knock-down lines, the expression is reduced to 34.66 ± 6.6. Interestingly, *OFP2* read count in over-expression line was found to be reduced to 2.6 ± 2.03 indicating a strong feed-back mechanism that limits *OFP2* in over-expression lines. It is paradoxical as despite having highly reduced amount of *OFP2* expression, over-expression lines still exhibit a strong stunted growth phenotype as opposed to knock-down lines.

To identify the biological pathways possibly affected by *OFP2* to exert such strong and contrasting phenotype in knock-down and over-expression lines, differentially expressed genes were studied. Genes involved in hormone signaling, cell wall synthesis and expansion, stress response, organ development and important transcription factors were found to be significantly deregulated in both genotypes. To systematically connect these genes with observed phenotype, four important categories emerged most direct explain the phenotypes. Those four categories are: (i) cell wall expansion and general organ development genes, (ii) root architecture defining genes, (iii) hormone-responsive genes and (iv) dormancy and seed germination related genes (Figure 8E-H). Among all the genes, cell wall expansion genes constitute one of the most significantly enriched functional cluster. Severe down regulation of expansins in *oex-ofp2* and mild upregulation in *amiR-ofp2* directly explains the opposite cell size phenotype affecting growth through seedlings to mature plants. The α-expansin genes *EXPA7* (log2FC: −8.77x in *Oex*, +1.27x in *amiR*), *EXPA4* (−3.20x / +1.02x), *EXPA17* (−1.75x / +1.30x), *EXLA3* (−1.78 x/ +1.29x), *EXPA18* (−3.67 x/ −1.44x), and *EXPB1* (−5.46x / −0.72x) showed a clear reciprocal expression patterns between conditions. Expansins loosen the cell wall by disrupting non-covalent bonds between cellulose and xyloglucan, enabling turgor-driven expansion without enzymatic hydrolysis (Cosgrove, 2005). Another important set of gene are xyloglucan endotransglucosylase/hydrolases *XTH12* (log2FC: −9.68x in *Oex*, −2.04x in *amiR*), *XTH26* (−9.53x / −2.01x), and *XTH21* (−8.79x / +0.01x), and the cellulose synthase-like genes *CSLC12* (−8.37x / −1.62x), *CSLB5* (−8.04 x/ −1.68x), and *CSLB1* (−8.22x / −0.44x).

The XTH family enzymes remodel the cellulose–xyloglucan network by transglycosylation and hydrolysis, enabling controlled cell wall yielding during growth (Rose et al., 2002; Nishitani and Vissenberg, 2007). Additional deregulated members of the cell wall machinery from the full dataset include *CESA3, CESA6, COBRA, FLA3*, and multiple *XTH* and *EXT* family genes (Supplementary Table S1 & S2). Pectin methylesterase *PME3* (−1.05x / +2.59x) displayed one of the most important reciprocal pattern as its suppression in *oex-ofp2* stiffening the pectin matrix and its strong induction in *amiR-ofp2* softening it contributing to the contrasting organ size phenotypes.

Analysis of differentially regulated genes against root development gene ontology identified enrichment of genes mainly involved in polar auxin transport, cytokinin signaling, epidermal cell fate specification and radial tissue patterning. Among all the genes in this category, *SHY2/IAA3* (log2FC: −3.49x in *Oex*, +1.02x in *amiR*) showed the clearest reciprocal regulation. It is known for its central role in coupling auxin and cytokinin inputs to control cell differentiation at the root meristem transition zone (Dello Ioio et al., 2008). Among auxin efflux carriers *PIN1* (−0.51x / +0.34x), *PIN7* (−2.22x / −2.14x) and *PIN3* (+0.45x / +0.45x) are deregulated indicating disrupted polar auxin transport. Notably, *PIN7* is repressed in both, and *PIN3* is slightly upregulated in both, showing indirect network effect for these genes. *GH3.17* (−1.45x / −0.45x), an IAA-amido synthetase controlling local auxin inactivation, was more strongly suppressed in *oex-OFP2*, and cytokinin receptor *AHK3* (−0.51x / +0.24x) showed mild reciprocal regulation together indicating that both auxin and cytokinin signalling arms of the meristem-to-elongation transition are differentially modulated. Additional deregulated genes from this sub-module including *ARR1, ARR12, AHP6, LAX3*, and *AXR2* are provided in Supplementary Table S1 and S2. *WER* (−9.26x / −0.08x) is one of the most strongly suppressed gene in the *oex-ofp2* transcriptome, with essentially no change in *amiR-ofp2*. It is a R2R3-MYB master regulator of non-hair (N-position) cell fate controlling the MBW transcription factor complex together with GL3 (−3.29x / −0.60x), EGL3 (−3.22x / −0.14x), and TTG1 to activate *GL2* (−1.61x / +0.01x) and repress root hair differentiation in N-position cells (Lee and Schiefelbein, 1999; Bernhardt et al., 2003). This deregulated complex seems to disturb epidermal cell identity explaining altered root hair density and patterning. The position-sensing receptor kinase *SCM* (−1.19x / +0.05x) and the lateral inhibitor *CPC* (−0.95x / −0.07x) were also suppressed in *oex-OFP2*, indicating disruption at multiple levels of the patterning hierarchy (Shibata and Sugimoto, 2019). In the radial patterning sub-module, *SHR, SCR*, and *PHB* were unchanged in both conditions, indicating that ground tissue specification is not a primary target of *AtOFP2*; however, *RBR1* (−0.53x / +0.17x) showed mild reciprocal regulation, suggesting a subtle modulation of the proliferation-to-differentiation balance at the quiescent centre (Cederholm et al, 2012).

Plants being immobile, phytohormones are the main chemical messengers controlling and coordinating all internal processes. Transcriptome data showed deregulation of hormone responsive genes for auxin, cytokinin, GA and brassinosteroid and ABA pathways, but not in a reciprocal way among both genotypes indicating a complex hormone signalling in these genotypes. For auxin, along with PIN transporters other important deregulated genes were YUCCA biosynthesis genes like *YUC3* (−2.90x / −0.56x) and *YUC9* (−1.91x / +0.04x) indicating reduced local IAA production. Important cytokinin pathway genes deregulated in both genotypes are Histidine containing phosphotranfer protein *AHP2* (1.12x / −0.02x) which was suppressed only in *oex-ofp2*, while the type-A negative feedback regulator *ARR7* (−2.73x / +0.64x) showed reciprocal regulation indicating attenuated cytokinin signaling in over-expression and enhancement in knock-down lines (Tran and Ruszkowski, 2025). For gibberellin signalling, *GA2OX2* (+1.72x / +0.55x) encoding GA-deactivating 2-oxidase was induced in both, but more strongly in *oex-ofp2*, while the DELLA repressor *RGL1* (+1.06x / −0.27x) showed a clear reciprocal pattern. An important GRAS intermediate *SCL3* (+1.62x / +0.25x) was induced in both, but strongly in *oex-ofp2*. Brassinosteroid signaling genes are mainly deregulated in over-expression lines and mostly unaffected in amir lines. *BSE1-Int* (−1.92x / +0.05x) an important BR output TF is suppressed in *oex-ofp2* while its upstream stimulators *BEE1* (+4.88x / n.d) and *BEE3* (+3.83x / n.d) upregulated in *oex-OFP2* with no change in *amiR-OFP2*. This dissociation between BR-stimulatory transcription factors and their canonical downstream output indicates a non-functional BR signalling state in *oex-OFP2* roots (Friedrichsen et al., 2002).

Another important set of deregulated genes revealing the reciprocal phenotypes for seed dormancy and germination are ABA signaling related genes. Abscissic acid is the most important phytohormone impacting seed biology. Most severely deregulated gene in this category is *MYB61* (−8.61x / −0.36x), a critical regulator of seed coat testa differentiation (Penfield et al., 2001). Three important attenuators of ABA signalling significantly suppressed in *oex-ofp2* are *HAI2* (−4.35x / +1.29x), clade A PP2C phosphatase that inactivates SnRK2 kinases and weakens ABA responses (Bhaskara et al., 2012); *PUB19* (−2.21x / +2.76x), the U-box E3 ligase causing ABA hypersensitivity in mutants (Liu et al., 2011); and *MYB96* (−2.02x / +1.28x) and *MYB41* (−1.52x / +1.31x), which regulate ABA-dependent seed coat wax and suberin deposition respectively (Seo et al., 2011; Kosma et al., 2014). Primary ABA 8′-hydroxylase catabolosing ABA in dry seeds is also suppressed in both genotypes but severely in *oex-ofp2* explaining accumulation of ABA in dry seeds enhancing dormancy (Okamoto et al., 2006).

### Validation of transcriptome data

To draw meaningful conclusions from transcriptome data its reliability and reproducibility check was necessary. The reliability of transcriptome data was validated by randomly selecting few candidate genes for expression analysis using quantitative real-time RT-PCR. For transcriptome data RNA from roots of 7-day old seedlings was used. To ensure a valid comparison for validation of these genes RNA was isolated from roots of 7-day old seedlings of WT, *oex-ofp2* and *amiR-ofp2*. All genes followed the same trend in expression analysis as transcriptome data. Nearly 94% decrease in the transcript level of *AT4G34580* (*CAN OF WORMS1 / COW1*) in *amiR-ofp2* lines (log2FC = - 4.097x) and a 93.2% decrease in *oex-ofp2* lines (log2FC = -3.88x) as compared to WT. Two tailed t-test analysis indicated a statistically significant decrease in the transcript level of *AT4G34580* in *amiR-ofp2* lines (p-value= 0.048<0.05) and in *oex-ofp2* lines (p-value= 0.036<0.05). Similarly, statistically significant downregulation of *AT2G33790* (*AGP30*) with -5.026x and -4.81fold-change was observed in *amiR-ofp2* and *oex-ofp2* lines respectively as compared to WT. Nearly 99% down-regulation in the expression of *AT1G04240* was observed in roots of *oex-ofp2* lines (log2FC= -6.64x; p-value= 0.0035<0.01), and 98.25% downregulation (log2FC= - 5.8366x; p-value= 0.0057<0.01) in *amiR-ofp2* as compared to WT. The transcript level of *AT5G58010* was found to be reduced by 49% (Log2FC= -0.97x; p-value= 0.39>0.05) in *amiR-ofp2* lines, and 76.7% (Log2FC= -2.1x; p-value= 0.13>0.05) in *oex-ofp2* as compared to WT (Figure 9).

**Figure 9:**
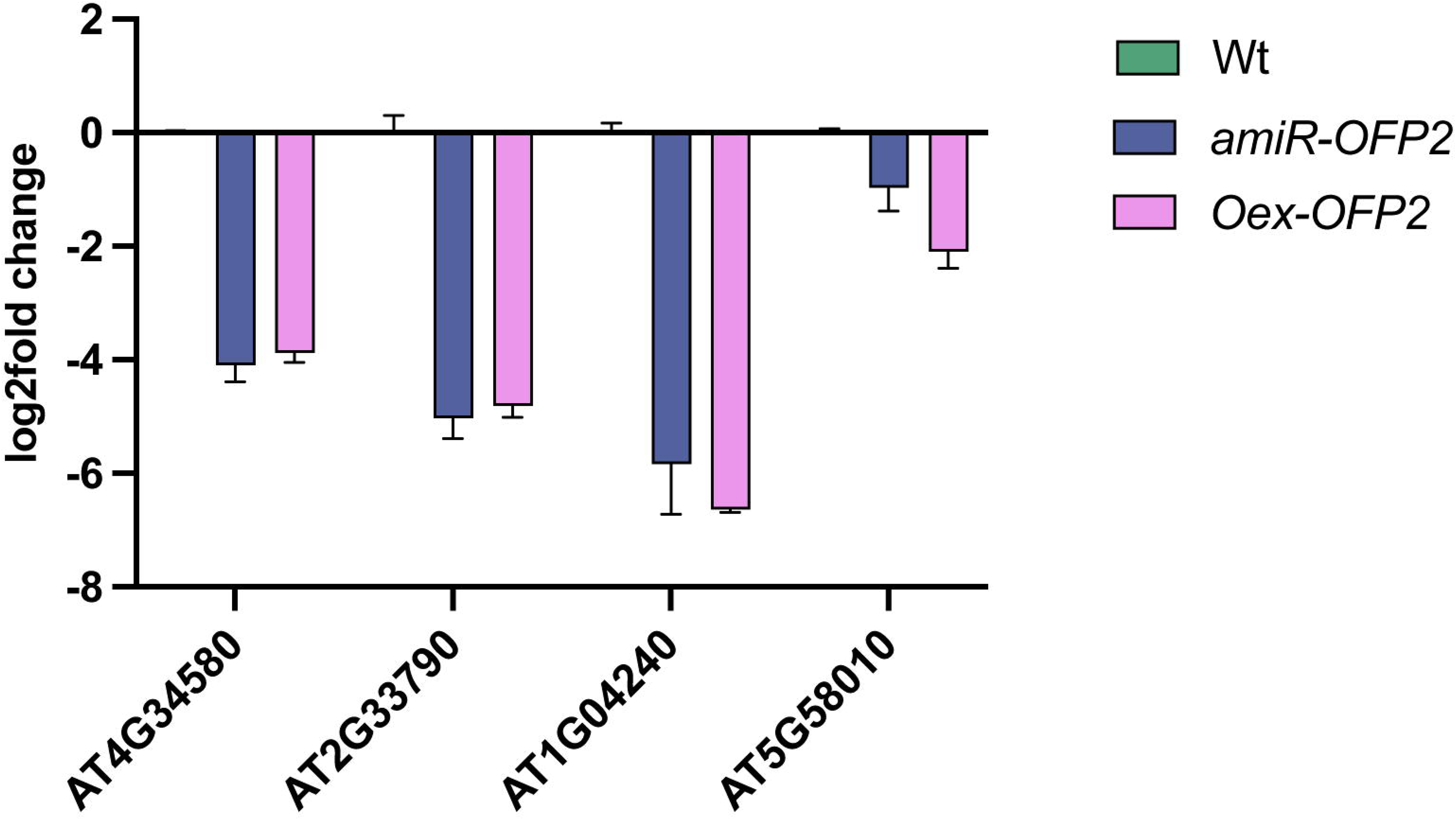
Validation of transcriptome data using qRT-PCR of (A) *AT4G34580*, (B) *AT2G33790*, (C) *AT1G04240*, and (D) *AT5G58010*. “*” and “**” represents statistically significant differences at (p<0.05), and (p<0.01), respectively; ns= non-significant.

**Figure 10:**
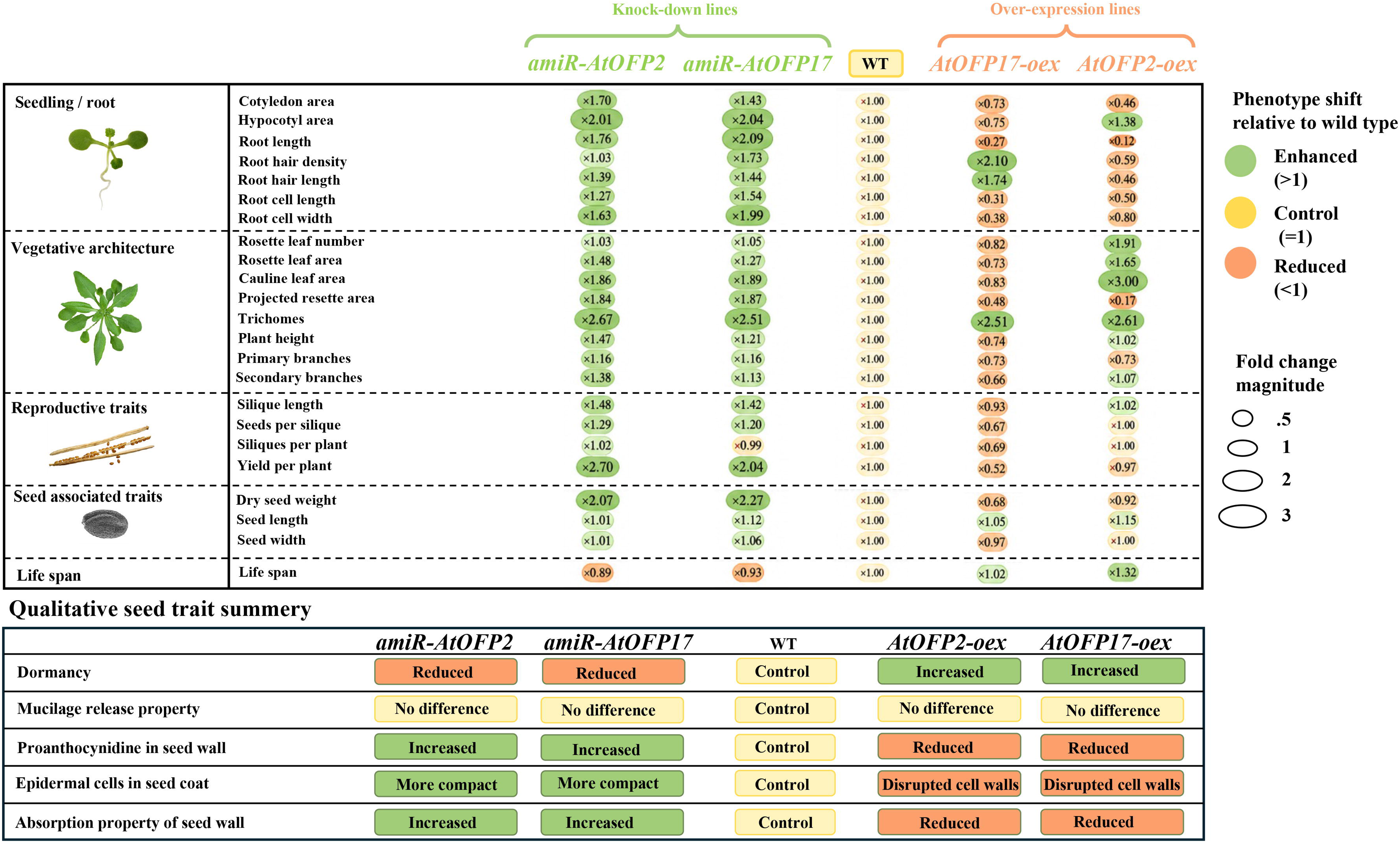
Compilation of traits measured in *amiR-AtOFP2, amiR-AtOFP17, WT, oex-AtOFP2,* and *oex-AtOFP17* mutants. Green color represent enhanced traits, orange color represent reduced traits and yellow color represent traits in control plants and statistically non-significant change.

## Discussion

Ovate Family Proteins (OFPs) are a class of plant-specific transcription regulators, unified by a highly conserved C-terminal OVATE domain. First identified in tomato (*Solanum lycopersicum*) for their role in regulating fruit shape, the OVATE locus encodes a nuclear-localized transcriptional repressor and shown to modulate key developmental and architectural traits. Based upon full and partial structure of ovate domain, OFPs have been divided into Ovate-OFPs and Ovate-Like OFPs (Chahar et al. 2021). The OFP gene family originated and expanded early in Bryophyta, encompassing protein variants with both complete and partial OVATE domains (Chahar et al. 2023). This early diversification suggests an adaptive advantage for functional specialization. Ancestral forms of specific *Arabidopsis* OFPs, such as *AtOFP2* and *AtOFP17*, are hypothesized to have evolved in the last common ancestors of Embryophyta (land plants), being notably absent from Rhodophyta and Chlorophyta, and points to their emergence as a key innovation in the evolution of land plants. A unique head-to-head gene organization for certain OFP pairs (e.g., *AtOFP2-AtOFP17* on chromosome 2 and *AtOFP4-AtOFP20* on chromosome 1 in *A. thaliana*) is observed exclusively in Spermatophyta (gymnosperms and angiosperms), with a duplication event in core Brassicaceae approximately 32-54 million years ago (Chahar et al. 2021, 2023). This genomic arrangement suggests potential co-regulation or specialized functional interplay between these gene pairs. Understanding domain integrity, its impact on molecular function and final phenotypic outcome is the central theme of this study as compete domain might confer full repressive activity or specific interaction capabilities, while a partial domain could lead to altered binding, modified interactions or even a dominant-negative effect.

To date, all reports on functional characterization have been limited to OFPs with complete Ovate-domain (Ovate-OFP), and characterization of Ovate-Like OFPs has been lacking leaving impact of domain intergrity on functional outcomes unresolved. We therefore performed comparative functional characterization of Ovate-OFP (*AtOFP2*) and Ovate-Like OFP (*AtOFP17*) that are arranged in head-to-head manner, by generating artificial microRNA-mediated knock-down, loss-of-function mutants (Schwab et al. 2005, 2006) and gain-of-function mutants (Zhao et al., 2017). To understand their comparative impact on plant phenotypes we studied 28 phenotypic characters in mutant plants. Starting from seed germination to final yield of plants a through comparison of each trait provided significant information to hypothesize that these two paralogs operate majorly in similar way as repressors, but their individual nuanced effects indicate sub-functionalization.

At juvenile stage, six traits- cotyledon area, hypocotyl area, root length, root hair density, root hair length and root cell dimensions were quantified and comparative analysis revealed that *AtOFP2* and *AtOFP17* exert repression in distinct ways, with *AtOFP2* showing stronger effects on some traits while *AtOFP17* severely impacting others. Overall, both genes showed repressor effects with over-expression mutants being smaller and knock-down mutant being larger than wild type due to release of transcriptional repression. For cotyledon area and primary root length, repression was most pronounced in *oex-ofp2* lines indicating *AtOFP2* as a stronger repressor of organ initiation and root elongation. However, hypocotyl growth was more severely repressed in *oex-ofp17*, pointing to *AtOFP17* as a stronger regulator of shoot elongation. Root hair traits also highlighted a divergence: *AtOFP2* acted as a suppressor, reducing hair density and length, whereas *AtOFP17* showed the opposite trend, with *oex-ofp17* lines displaying increased density and longer hairs, suggesting an antagonistic or modulatory role. Finally, in root cell dimensions, *AtOFP17* overexpression produced extreme reductions compared to *AtOFP2*, underscoring *AtOFP17*s stronger effect on cellular architecture. Also, ∼1.5 fold larger cells in *amirRNA* lines and ∼0.5 fold smaller cells in *oex* lines indicate towards variation in cell size, and not cell number being the main reason of observed differences in total plant growth patterns. Collectively, these data indicate that along with a general cell size repression that both genes exert, *AtOFP2* functions as a broad-spectrum repressor of cotyledon expansion and root elongation, while *AtOFP17* exerts more severe repression of hypocotyl growth and root cell morphology, and simultaneously enhances root hair traits. Among all the traits analyzed, primary root was most strikingly affected in both genes and in a similar way. Further, we complemented phenotypic data with transcriptome data of knock-down and over-expression mutants of *OFP2* from root tissue of 7-day old seedling which provided mechanistic framework on molecular mechanism to interpret observed phenotypes. From transcriptome data, deregulation of cell wall expansion gene sets provides a most direct explanation for altered cell and organ size. Multiple expansin family genes like *EXPA7, EXPA4, EXPA18, EXPB1, EXPA17, EXLA3*; Xyloglucan endotransglucosylase/hydrolases *XTH12*, *XTH26*, and *XTH21;* and cellulose synthase-like genes *CSLC12*, *CSLB5*, *CSLB1* are down-regulated in *oex-ofp2* mutants and are up-regulated in *amiR-ofp2* mutants (figure 8E). In Arabidopsis, cell size is determined by a controlled balance between cell wall expansion and structural reinforcement (Czesnick and Lenhard, 2015). Expansins are non-enzymatic wall-loosening proteins which induce turgor-driven cell expansion by disrupting the load-bearing hydrogen bonds between cellulose and hemicellulose (Cosgrove, 2005). Cellulose synthases construct the highly rigid cellulose microfibrils at the plasma membrane. And XTH enzymes modify the primary cell wall’s hemicellulose network. They temporarily cut xyloglucan tethers and reconnect them, allowing the cellulose framework to physically relax so cells can expand (Rose et al., 2002; Nishitani and Vissenberg, 2007). *XTH21* in particular is required for primary root growth by regulating cellulose deposition (Liu et al., 2007). Pectin methylesterase *PME3* and cellulose synthase-like genes *CSLC12*, *CSLB5*, *CSLB1* were also downregulated, indicating broad suppression of the cell wall biosynthetic and remodelling machinery. All these genes are up-regulated in *amiR-ofp2* mutants explaining opposite phenotype in these lines. *AtOFP2* has been shown to interact with TONNEAU2 to regulate cortical microtubule rearrangement for cell size regulation. This interaction is regulated by Brassinosteroid signalling (Zhang et al, 2020). Two BR-responsive bHLH factors *BEE1* and *BEE3* and one BR-activated transcription factor *BES1* are also downregulated, indicating that *AtOFP2* suppresses BR-mediated cell elongation pathways. Triple knockout of *BEE1, BEE2* and *BEE3* displays stunted growth phenotypes similar to *oex-ofp2* (Friedrichsen et al., 2002).

Severe retardation of roots in *oex-ofp2* more than other tissues can be explained by disturbed epidermal and radial patterning of roots which can be explained by down-regulation of genes involved in these processes. Analysis of transcriptome data showed downregulation of *WER* (FC -9.26x), *GL3* (-3.29x), *EGL3* (-3.22x), *GL2* (-1.61x), *CPC* (-0.95x), and *SCM* (-1.19x), components of the MYB-bHLH-WD40 complex that determines non-hair cell fate. In non-hair cells, WER forms a trimeric complex with GL3/EGL3 and TTG1 to activate *GL2* transcription, thereby repressing root hair cell differentiation (Lee and Schiefelbein, 1999; Bernhardt et al., 2003, 2005). The small R3 MYB repressor CPC moves from non-hair to hair cells and competes with WER for GL3/EGL3 binding, promoting hair cell fate; CPC transcription itself is activated by the MBW complex in non-hair cells (Wada et al., 2002; Kurata et al., 2005). RSL2 and RSL4, two important downstream genes in this pathway are also severely down-regulated by log2 fold value of -10.1x and -9.6x respectively representing an amplified effect of cascade from top to bottom in a pathway. SCRAMBLED (SCM), an atypical leucine-rich repeat receptor-like kinase which is also down-regulated in *oex-ofp2*, initiates position-dependent patterning by modulating WER accumulation (Kwak et al., 2005). SCM itself works under the transcriptional feedback regulation of MBW complex (Kwak and Schiefelbein, 2007). Suppression of this entire module in *AtOFP2-oex* seem to disrupt the N-cell/H-cell boundary and alter epidermal patterning. Other important gene down-regulated in *oex-ofp2* is *RBR1* (RETINOBLASTOMA-RELATED 1), which coordinates cell cycle exit with differentiation in the root meristematic zone and interacts directly with SCR (Wildwater et al., 2005; Cruz-Ramírez et al., 2012), further suggesting broad disruption of the proliferation-to-differentiation transition under *AtOFP2* overexpression.

The dysregulation of several hormone-responsive genes in over-expression mutants suggests that the widespread repressive action of OFP2 can be achieved by a coordinated multi-hormone signaling cascade. Central node regulator of this network is *SHY2/IAA3* (log2FC: −3.49x in *Oex*; +1.02x in *amiR*) which is reciprocally expressed in both genotypes also validated by RT-PCR. In multicellular organisms, organ size is determined by the balance between cell division and differentiation. In Arabidopsis roots, this balance is mediated by auxin and cytokinin using a regulatory circuit converging on *SHY2* gene. It is a primary integrating node of the auxin–cytokinin antagonism governing root meristem size. In wild-type roots, cytokinin activates *SHY2* via ARR1-mediated direct promoter binding, causing *SHY2* to repress *PIN* auxin efflux carriers and drive transition-zone differentiation; auxin conversely promotes SHY2 degradation to sustain proximal meristem cell division (Ioio et al., 2007, 2008). In the cytokinin relay, both His-phosphotransfer protein *AHP2* (−1.12x in *Oex*; −0.02x in *amiR*) and type-A response regulator *ARR7* (−2.73x in *Oex*; +0.64x in *amiR*) was suppressed in *oex-ofp2*, impairing cytokinin signal transduction (Hutchison et al., 2006). The concurrent suppression of both *AHP2* and *ARR7* creates an incoherent feed-forward state in which cytokinin signalling is neither properly transmitted nor appropriately attenuated. *BES1* (−1.92x in *Oex*; +0.05x in *amiR*) directly binds the promoters of both *PIN7* and *SHY2* to promote meristem expansion, and its loss restricts root elongation (Li et al., 2020). The simultaneous suppression of *BES* and *SHY2* therefore dismantles the three-hormone BR–auxin–CK equilibrium required for optimal root growth.

In mature plants, just like seedling stage, loss of function of both *AtOFP2* and *AtOFP17* promoted overall growth of plants which was seen in expanded leaf area, taller plants, a greater number of branches, more and bigger siliques and total plant yield. This further confirms that both genes work as repressors of plant architecture. However, over-expression revealed a striking difference in their functionality dictated by the mutants after being transferred in pots. *OFP17* showed a classic repression in plants through all the developmental stages with knock-down plants being bigger with more yield and over-expression being smaller with lesser yield. However, *oex-ofp2* plants showed constant degeneration of true leaves after transferring to pots (Figure 3G), maintaining cotyledonary leaves only, for a prolonged period. Degeneration of newly emerging leaves seems like primordium viability failure causing a developmental arrest. This can be explained by the suppression of PIN1 (-0.51x) which depletes auxin transport machinery and YUCCA flavin monooxygenases *YUC3* (−2.90x) and *YUC9* (−1.91x) which catalyse the rate-limiting IPyA-to-IAA step in auxin biosynthesis depleting the auxin maximum required for meristem maintenance (Yadav et al. 2023). Once this initial repression was overcome, *OFP2* over-expression plant grew normally similar to wild type plants and no further signs of repression were seen till they were finally harvested. This suggests that the initial over-expression driven repression was not constitutively sustained throughout development. This pattern of early lethality evidenced by only 10% survived *oex-ofp2* mutants; severe growth repression and then recovery hints towards dosage sensitive transcriptional regulation of *OFP2*. Surviving plants represent individuals in which this repression might have activate a feedback loop mechanism which is a common for transcription factors, or post translation gene silencing which is also common for 35S-driven high-copy transgenes in *Arabidopsis* (Fagard and Vaucheret, 2000; Matzke and Birchler, 2005). A lot of transcription factors are regulated through feedback loop mechanism like WUS-CLV3 (Schoof et al, 2000); miRNA156-SPL (Wu et al, 2009); TFL1-LFY (Huang et al, 2026) to modulate plant growth and specific developmental phases.

Nevertheless, the persistence of an extreme short-root phenotype in these same plants indicates that sufficient stable OFP2 protein remained to exert targeted repression. This strong repressive phase at juvenile stage caused a pronounced heterochronic shift in *oex-ofp2* mutants as they matured equal to wild type plants but that took extra 5-weeks extending their life span to 15-16 weeks compared 12-13 weeks for wild type and *amiR* plants. Heterochrony is regulated by the conserved miR156/miR172-SPL pathway (Chuck et al, 2007; Wu et al, 2009). Presence of SQUAMOSA PROMOTER BINDING site in promoter of OFP2 supports potential activation of this regulatory pathway.

Seed associated phenotypes showed a consistent reciprocal pattern for both genes. Though *ofp2* suppression was not seen in mature plants but it was similar to *ofp17* for seed physiology. Knock-down of both genes showed reduced dormancy (seed germination within 12 hours), high proanthocyanidin levels (dark purple seeds), compact epidermal seed coat cells with high absorbance properties. On the contrary, over-expression mutants were dormant (started germination around 48 hours), accumulated low proanthocyanidin levels (only at chalazal ends), distorted epidermal cells with less absorption properties while mucilage release was unaffected across all genotypes, disassociating it from germination phenotypes. High PA levels are associated with dormancy but here paradoxically, amiR mutants were less dormant with high PA levels. This can be because of the high permeability of seed coats. Transition from dormancy to seed germination is a highly coordinated process orchestrated by hormone signaling, chromatin remodeling, transcription reprogramming and epigenetic regulation. Central to this process is the balance between abscisic acid (ABA), which enforces dormancy, and gibberellic acid (GA), which promotes germination (Holdsworth et al., 2008; Graeber et al., 2012). Transcriptome data provided key molecular candidates for dormancy and germination which were considerably deregulated in *OFP2* mutants. *CYP707A1* (log2FC: −1.35x in oex-*ofp2*), encodes the key ABA 8′-hydroxylase responsible for ABA catabolism in dry seeds. Its suppression elevates ABA levels and enforces deeper dormancy, as *cyp707a1* mutants accumulate ABA to approximately 20-fold higher levels in dry seeds (Okamoto et al. 2006). *HAI2* (AT1G07430; log2FC: −4.35x in *oex-ofp2*), a clade A PP2C phosphatase that negatively regulates ABA signaling by inactivating SnRK2 kinases, was significantly downregulated. Reduced HAI2 activity would amplify ABA signalling and reinforce dormancy (Bhaskara et al., 2012). *PUB19* (AT1G60190), a U-box E3 ubiquitin ligase that negatively regulates ABA responses and whose down-regulation leads to ABA hypersensitivity and enhanced stomatal closure (Liu et al., 2011), was also downregulated (log2FC: −2.21x), further supporting a pro-dormancy ABA state in overexpression lines. *DAG2*, a *DOF*-type zinc finger transcription factor, positively regulates germination in part by activating *ECT1*, which promotes radicle emergence. Consistently, *dag2* mutants display reduced germination, requiring additional light or cold treatment and showing decreased sensitivity to GA (Gualberti et al., 2002; Li et al., 2024). Another downstream effector of GA is *ATMAN7*, which encodes a mannan endo-1,4-β-mannosidase that hydrolyzes cell wall mannans in the micropylar endosperm, enabling tissue loosening, seed coat rupture, and radicle protrusion; *atman7* mutants germinate more slowly (Fernandez et al., 2013). Finally, *BME3 (BLUE MICROPYLAR END 3*), a GATA-type zinc finger transcription factor expressed specifically in the embryonic axis, acts as a positive regulator of germination. Loss-of-function *bme3* mutants exhibit reduced germination rates even after stratification, underscoring its essential role in radicle emergence (Liu et al., 2005). Other important seed coat structure gene severely downregulated in *oex-ofp2* is *MYB61* (AT1G09540; log2FC: −8.61x). *MYB61* is a critical regulator of seed coat testa differentiation and mucilage deposition, and its loss impairs testa epidermal maturation and germination under stress conditions (Penfield et al., 2001). *MYB96* (AT5G62470; (log2FC: −2.02x), an ABA-responsive R2R3-MYB transcription factor that directly activates cuticular wax biosynthesis genes suggesting altered surface permeability of the seed coat (Seo et al., 2011). Distorted radial walls of epidermal seed coat cells, loose microfibrillar structure of seed walls in *oex-ofp2* can be a function of *MYB41* (AT4G28110; (log2FC: −1.52x), which activates ectopic suberin synthesis in response to drought and ABA, pointing to reduced suberization of seed coat cell walls (Cominelli et al., 2008; Kosma et al., 2014). Conversely, *EXPA5* (AT3G29030) was strongly upregulated (log2FC: +7.82x); expansin-mediated wall loosening in the micropylar endosperm is known to facilitate seed coat rupture and radicle protrusion (Cosgrove, 2005; Cho and Cosgrove, 2002), and its induction may represent a compensatory mechanism promoting germination competence. Together, concurrent supression of ABA catabolism along with its negative signalling regulation, distorted seed coat structure created a strong dormancy state which is further reinforced by GA and BR downregulation. Both these hormones antagonize ABA to promote germination (Holdsworth et al., 2008; Graeber et al., 2012; Friedrichsen et al., 2002).

## Conclusion

This study established that both *AtOFP2* and *AtOFP17* despite having difference in their domain integrity, works as general repressors of plant growth and development. Release of transcriptional repression of both genes yielded similar phenotypes in knock-down mutants revealing broadly same mode of action as repressors. However, distinct effects of over-expression mutants indicated towards sub-functionalization as a function of domain structure. Strong repression of *AtOFP2* at juvenile stage, death of 90% over-expression mutants and final recovery to achieve growth comparable to wild type plants indicate that *AtOFP2* is a very strong transcriptional repressor possibly controlled by feedback-loop mechanism. However, exact mechanism for its function and control needs to be evaluated. Due to partial deletion of ovate domain, *AtOFP17* seems to have milder repression which can be tolerated by plants throughout entire life cycle with compromised growth and yield. Transcriptome data analysis indicates important and complex position of *AtOFP2* which seems like operating at a crosstalk node of phytohormones. Deregulated genes related to auxin (*YUC3/9–SHY2–PIN1/3/7–GH3.17*), cytokinin (*AHK3–ARR1/12*), GA (*GA20ox2–RGL1–SCL3*), BR (*BES1–BEE1/3*), and ABA (*CYP707A1–HAI2–PUB19*) along with downstream consequences for cell wall remodelling (*EXPAs, XTHs, CSLs*), epidermal patterning (*WER–GL3/EGL3–TTG1–GL2–CPC–SCM*), radial tissue identity (*SHR–SCR–RBR1*), and seed coat integrity (*MYB61, MYB96, MYB41, EXPA5*) hints towards a higher position of *AtOFP2* in hierarchy of gene regulatory networks. Most importantly, higher yield in knock-down mutants makes them attractive targets for crop improvement. Future work dissecting the protein–protein interactions mediated by the complete versus partial OVATE domain, and identifying their direct transcriptional targets, will be critical to fully resolving the mechanistic basis of their functional specialization.

## Acknowledgments

SD would like to acknowledge Institute of Eminence (Ref. No./IoE/2023-24/12/FRP, Ref. No./IoE/2024-25/12/FRP and Ref. No./IoE/2025-26/12/FRP), and University of Delhi for Infrastructure and financial support. The award of JRF/SRF from CSIR to NC and SY, from UGC to EP, and DBT-BioCare to MD is gratefully acknowledged.

## Declarations

### Ethical Approval

Not applicable

### Authors’ contributions

NC and SD designed the study. NC generated all biological material, phenotyping, imaging and performed all analysis. EP and SY helped NC in generating transcriptome data and validation. BR helped NC in computational analysis of transcriptome data. MD helped in generation of reverse genetic mutants. NC and SD analysed all data and wrote the MS. SD was responsible for funds acquisition. All authors have read and agree with the findings.

### Availability of data and materials

All data generated in the study are submitted as supplementary files. The transcriptome data is available at SRA under project ID PRJNA1401268

## Supplementary data

**Supplementary Figure 1:** Genetic cassette of pGWB441 vector used for constitutive over-expression of *AtOFP2* and *AtOFP17* genes. [B] Line diagram of gene structure of *AtOFPs* cloned in pGWB441 vector. [C] Vector map of pGWB441

**Supplementary Figure 2:** [A], [B] Predicted stem-loop structure of *amiR-ofp2* precursor (ΔG: -72.4 kcal/mol) *and amiR-ofp17* precursor (ΔG: -76 kcal/mol); [C], [D] Gene structure of *AtOFP2* [C] and *AtOFP17* [D] showing site and sequence of amiRNA target and mature amiRNA.

**Supplementary Figure 3:** [A], [B] Schematic representation of strategy for engineering artificial microRNA in pRS300 backbone using Splicing Overlapping Extension (SOEing) PCR (Schwab, 2005). [C] Genetic cassette of artificial microRNA in pCHF3 vector. [D, E] Steady-state levels of target transcripts in knock-down and over expression lines detected through semi-quantitative RT-PCR analysis. Expression levels of Ubiquitin (UB), *AtOFP2* and *AtOFP17* in *amiR-OFP2/17*, WT and *AtOFP2/17*-oex cDNA from open flower.

**Supplementary Figure 4:** Comparison of cell length and width in roots from maturation zone of 15-days old seedling [A] DIC images of root cells from maturation zone of *amiR-ofp2*, WT and *oex-OFP2*. [B] Graphical representation of statistical comparisons of root cell length and width. [C] Statistical comparison of trichome density. [D] Trichome density on stem (left to right) of *amiR-ofp2*, WT and *oex-OFP2* plants. Bars represent mean ± SD, N=10, ns = no significant difference, *p=0.033, **p=0.02, ***p<0.001

**Supplementary Figure 5:** Comparison of cell length and width in roots from maturation zone 15-days old seedling [A] DIC images of root cells from maturation zone of *amiR-ofp2*, WT and *oex-OFP2* . [B] Graphical representation of statistical comparisons of root cell length and width. [C] Statistical comparison of trichome density. [D] Trichome density on stem (left to right) of *amiR-ofp17*, WT and *oex-OFP2* plants. Bars represent mean ± SD, N=10, ns = no significant difference, *p=0.033, **p=0.02, ***p<0.001

**Supplementary Figure 6:** Components of hormonal signalling pathway affected in *AtOFP2* mutants

**Supplementary Figure 7:** Components of circadian rhythm pathway affected in *AtOFP2* mutants

**Supplementary table 1:** List of primers used in this study

**Supplementary file 2:** Codes and parameters used in RNA-Seq analysis pipeline.

**Supplementary tables 2-20:** Raw data of phenotypic measurements of *AtOFP2* and *AtOFP17* mutants and their statistical analysis.

**Supplementary Table S21:** Lists of deregulated genes, significantly up and down regulated genes and transcription factors in over-expression genotype of *AtOFP2*.

**Supplementary Table 22:** Lists of deregulated genes, significantly up and down regulated genes and transcription factors in knock-down genotype of *AtOFP2*.

**Supplementary Table 23:** Read counts of transcripts among over-expression and knock-down genotype of *AtOFP2*.

