## Supplementary figures for "Head-to-head organized segmental paralogs *AtOFP2* and *AtOFP17* exhibit differential, spatio-temporal partitioning of function, and negative regulation of multiple developmental traits including seed-yield and root architecture"

### Supplementary figure 1

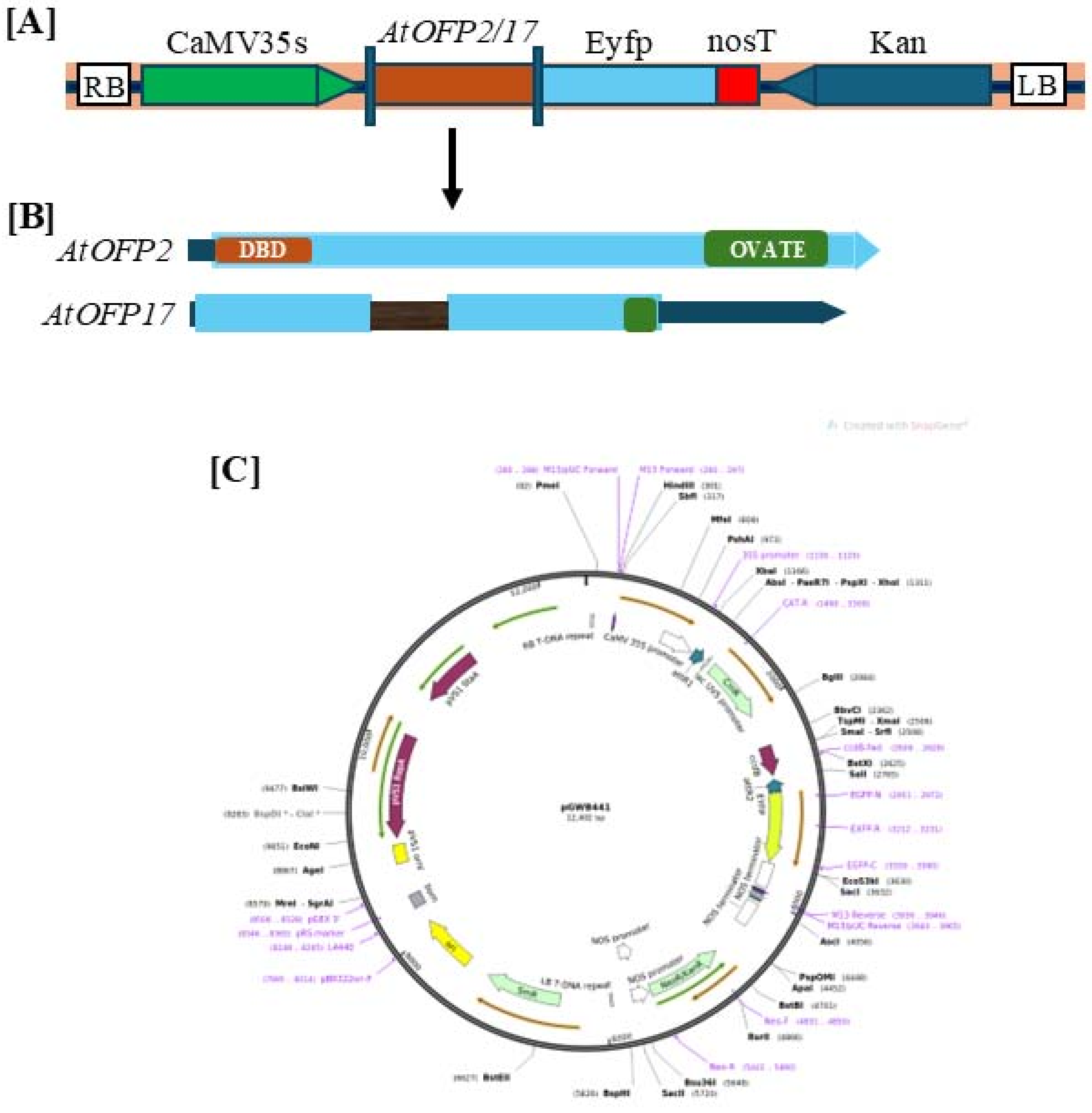

### Supplementary figure 2

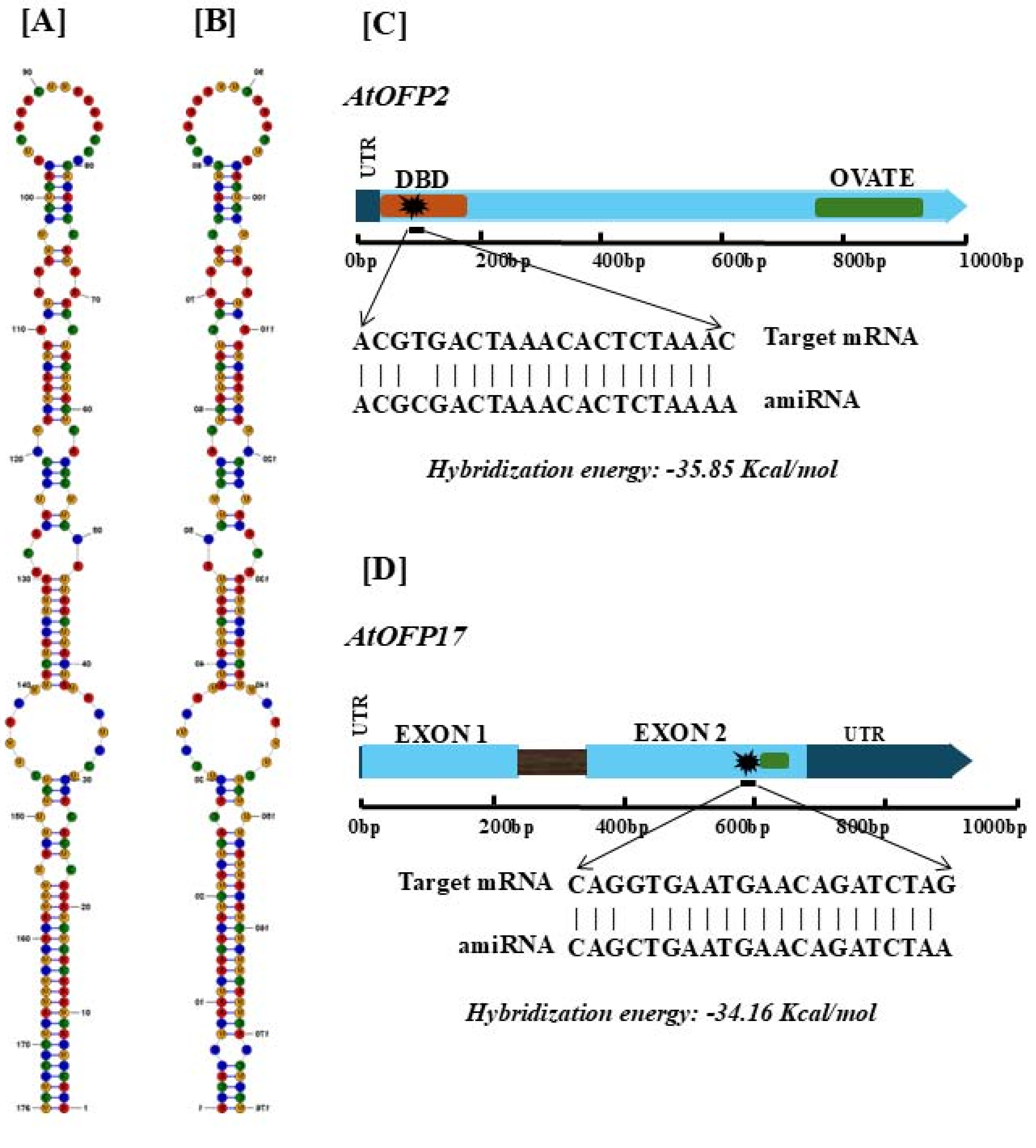

### Supplementary figure 3

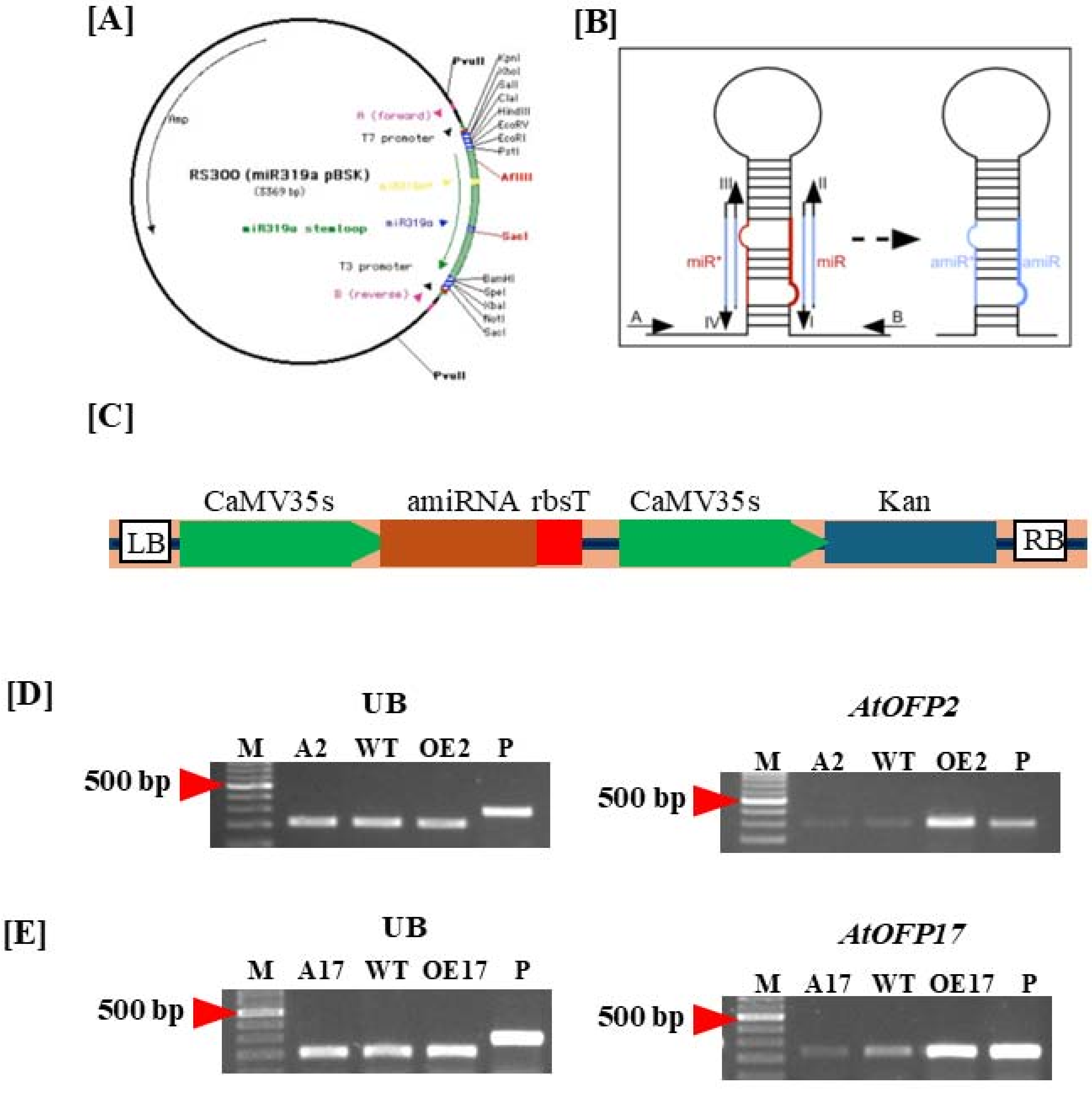

### Supplementary figure 4

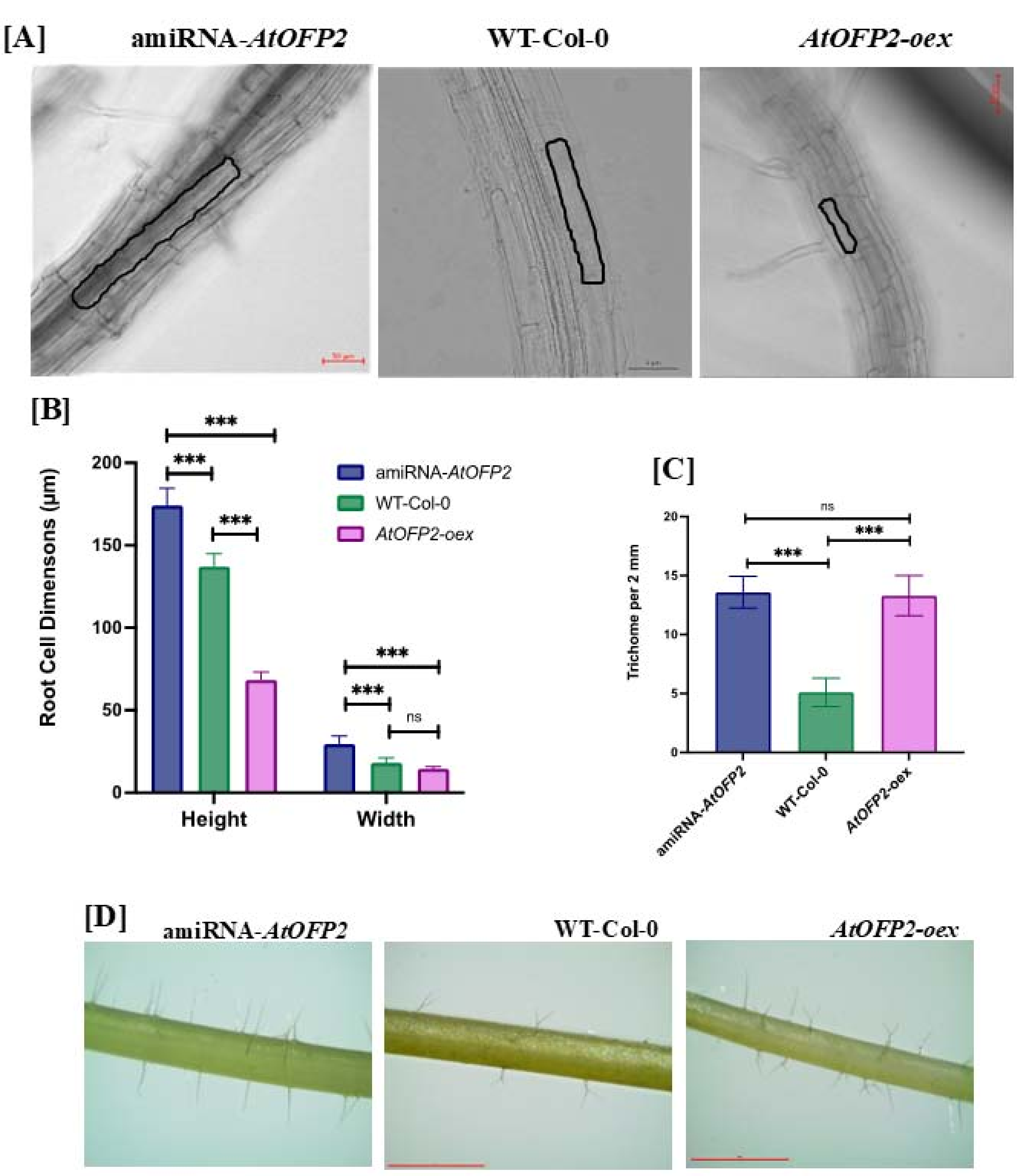

### Supplementary figure 5

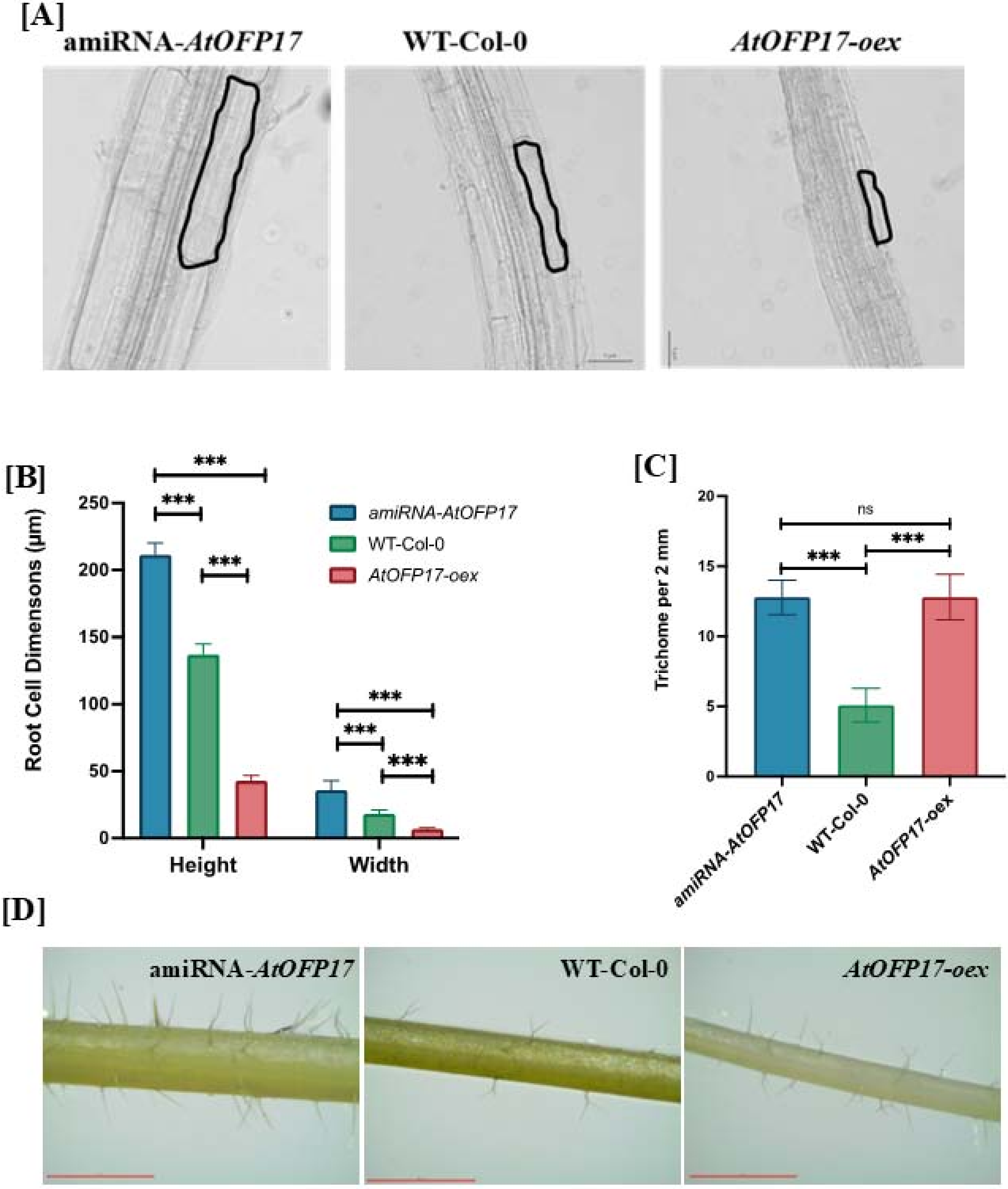

### Supplementary figure 6

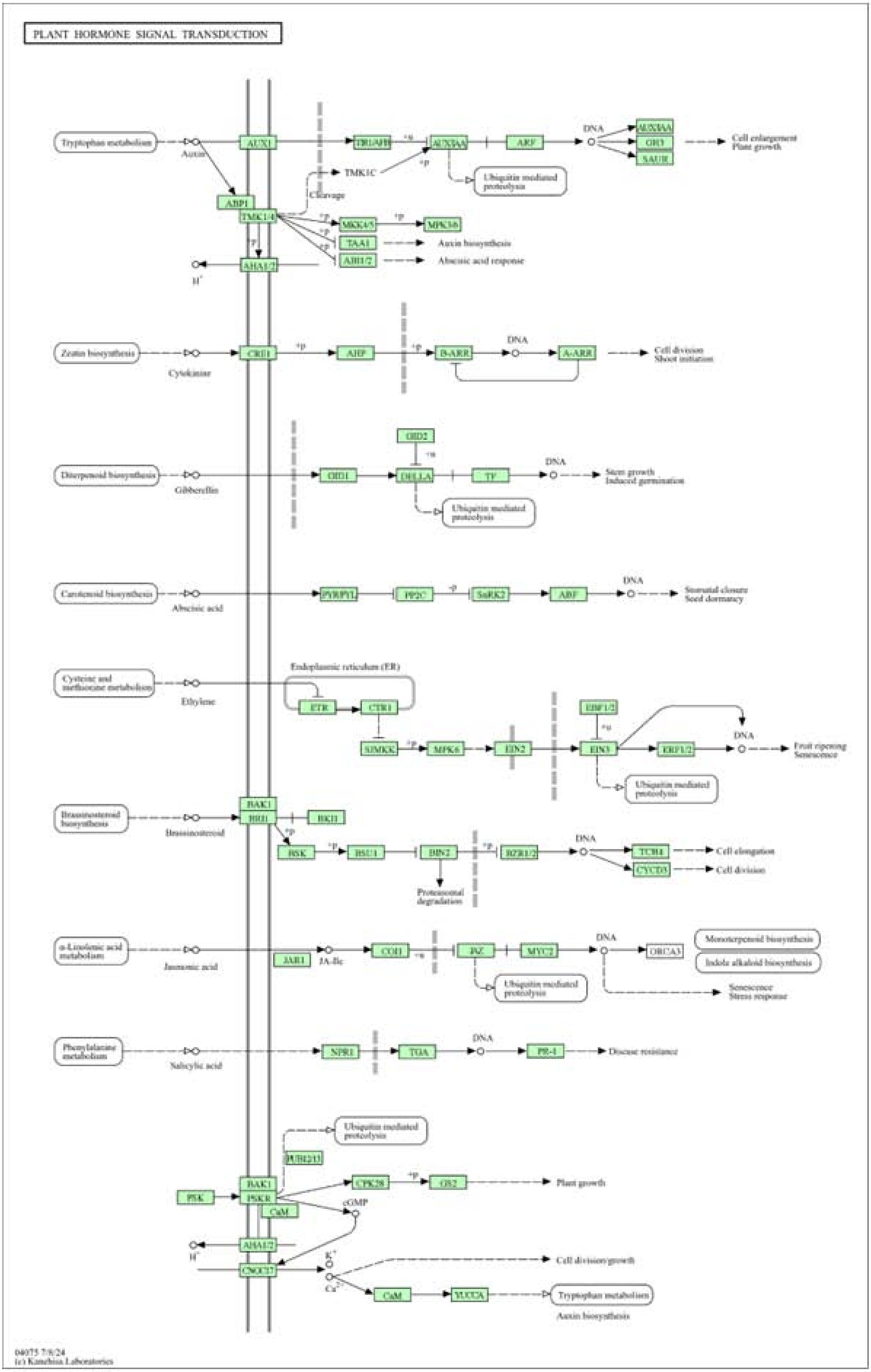

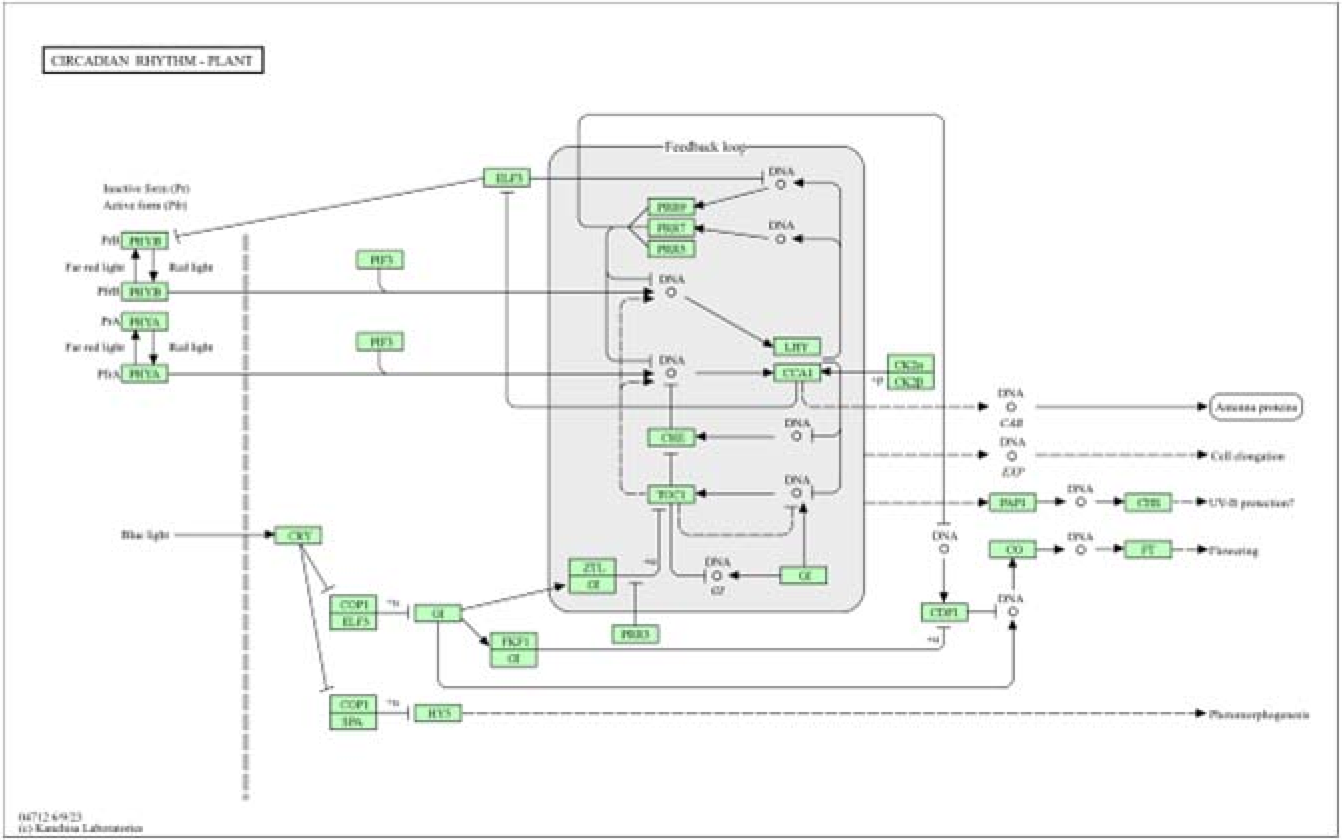
CIRCADIAN RHYTHM - PLANT
Feedback loop
Inactive form (Pr)
Active form (Pfr)
PHYB
PHYA
Far red light
Red light
Blue light
PIF3
ELF3
PRR9
PRR7
PRR5
DNA
LHY
CCA1
CK2α
CK2β
CHE
TOC1
ZTL
GI
PRR3
CRY
COP1
ELF3
FKF1
SPA
HY5
CDF1
CO
FT
PAP1
CHS
CAB
EXP
Antenna proteins
Cell elongation
UV-B protection?
Flowering
Photomorphogenesis
04712 6/9/23
(c) Kanehisa Laboratories
