## Supplementary file 1 for "Head-to-head organized segmental paralogs *AtOFP2* and *AtOFP17* exhibit differential, spatio-temporal partitioning of function, and negative regulation of multiple developmental traits including seed-yield and root architecture"

**Codes and parameters used in RNAseq analysis pipeline**

**Trimming of low quality reads and adapters in Trimmomatic** (v0.39):

trimmomatic PE -phred33 $paired_end_f1 $paired_end_f2 ILLUMINACLIP:TruSeq3-SE:2:30:10 LEADING:3 TRAILING:3 SLIDINGWINDOW:4:15 MINLEN:36

**Generation of reference genome index:**

STAR --runMode genomeGenerate --genomeDir TAIR_genome/ --genomeFastaFiles Arabi_TAIR10.1_genomic.fasta --sjdbGTFfile Arabi_genomic.gtf --sjdbGTFfeatureExon exon --genomeSAindexNbases 13

Alignment using STAR(2.7.10):

STAR --genomeDir TAIR_genome/ --outSAMattributes All --outSAMtype BAM SortedByCoordinate --quantMode GeneCounts --readFilesCommand zcat
--runThreadN 8 --outMultimapperOrder  Random --outWigType wiggle
--readFilesIn $f1 $f2 --outFileNamePrefix ${f1%%_1.fq.gz}
--sjdbGTFfeatureExon exon --sjdbGTFtagExonParentGene gene
--sjdbGTFfile Arabi_genomic.gtf --sjdbGTFtagExonParentTranscript transcript

**HTSeq-count** (v0.12.4):

htseq-count -f bam -m intersection-nonempty -t gene -i gene_id ${i} ../Arabi_genomic.gtf > $OUTFILE

**DESeq2 analysis in R**

data=read.csv('~/Downloads/nishu/ami_wt_count_table.csv',row.names=1)

col_ordering=c(1,2,3,4,5,6)
expMatrix=data[,col_ordering]

expMatrix=round(expMatrix)

expMatrix=expMatrix[rowSums(cpm(expMatrix)>1)>=2,]

#cpm=cpm(expMatrix)

#keep_genes=rowSums(cpm(expMatrix)>1)>=2

conditions=factor(c(rep('ami',3),rep('wt',3)))

exp_study=DGEList(counts=expMatrix,group=conditions)

exp_study=calcNormFactors(exp_study)

exp_study=estimateDisp(exp_study)

et=exactTest(exp_study,pair=c('ami','wt'))

topTags=topTags(et,n=NULL)

result_table=topTags$table

result_table=data.frame(sampleA='ami',sampleB='wt',result_table)

result_table$logFC=-1*result_table$logFC

write.table(result_table,file='ami_wt_DE_result.csv',sep=',',quote=F,row.names=T)

write.table(expMatrix,file='ami_wt_DE.count_matrix.csv',sep=',',quote=F, row.names=T)

-----------------------------------------
 
data=read.csv('~/Downloads/nishu/Oxp_wt_count_table.csv',row.names=1)

col_ordering=c(1,2,3,4,5,6)
expMatrix=data[,col_ordering]

expMatrix=round(expMatrix)

expMatrix=expMatrix[rowSums(cpm(expMatrix)>1)>=2,]

#cpm=cpm(expMatrix)

#keep_genes=rowSums(cpm(expMatrix)>1)>=2

conditions=factor(c(rep('oxp',3),rep('wt',3)))

exp_study=DGEList(counts=expMatrix,group=conditions)

exp_study=calcNormFactors(exp_study)

exp_study=estimateDisp(exp_study)

et=exactTest(exp_study,pair=c('oxp','wt'))

topTags=topTags(et,n=NULL)

result_table=topTags$table

result_table=data.frame(sampleA='oxp',sampleB='wt',result_table)

result_table$logFC=-1*result_table$logFC

write.table(result_table,file='Oxp_wt_DE_result.csv',sep=',',quote=F,row.names=T)

write.table(expMatrix,file='Oxp_wt_DE.count_matrix.csv',sep=',',quote=F, row.names=T)
